# Structural characterization of human endogenous retrovirus integration and strand transfer inhibition

**DOI:** 10.64898/2026.06.24.734183

**Authors:** Ane Barrena-Martin, Sandra Fuertes, Manuel Daza-Martin, Guillermo Abascal-Palacios

## Abstract

Human endogenous retroviruses (hERVs) are remnants of ancestral retroviral infections that have shaped the human genome through their capacity to mobilize and retrotranspose. Among these, hERV-K (HML-2) remains the most recently active family and its dysregulation is strongly associated with diverse cancers and neurodegenerative pathologies, yet the structural basis of its integration remains poorly understood. Here, we combine activity assays with high-resolution cryo-electron microscopy to resolve the hERV-K integration machinery in three distinct states: asymmetric target-DNA engagement, strand transfer, and pharmacological inhibition. Our structures reveal a compact architecture defined by a unique organization of the outer integrase domains, which distinguishes hERV-K from other known retroviral intasomes. Biochemical validation confirms the catalytic competence of this compact tetrameric assembly, which relies on specialized polar motifs to optimize synaptic stability while retaining sensitivity to competitive antagonism by strand transfer inhibitors. Notably, beyond canonical restriction by Raltegravir, we discovered that the drug binding stabilizes an unanticipated, “closed” conformation not observed in previously characterized intasomes. Together, these findings elucidate the molecular mechanism of endogenous retroviral integration and provide a structural framework for rational therapeutic targeting of hERV-K-driven diseases.

## Main

Transposable elements (TEs), which comprise retrotransposons and DNA transposons among others, are genetic sequences that drive genomic and functional diversity through their capacity to mobilize within the host genome^1^. Previously relegated to the status of “junk DNA”, TEs are now recognized as fundamental constituents of the human landscape, occupying nearly half of the human genome and acting as critical modulators of gene regulatory networks^2,3^ and disease susceptibility^4–6^.

Among retrotransposons, human endogenous retroviruses (hERVs) represent remnants of ancient retroviral infections that integrated into germline DNA millions of years ago and subsequently became inherited as stable genomic components^5,7,8^. Accounting for approximately 8% of the human genome, hERVs exhibit structural homology to exogenous retroviruses such as the Human Immunodeficiency Virus (HIV) and the Human T-lymphotropic Virus (HTLV), which are the causative agents of AIDS and adult T-cell leukaemia/lymphoma, respectively^9^. From a genetic point-of-view, these elements are flanked by Long Terminal Repeat (LTR) sequences, which serve as essential regulatory regions and harbor the promoters and enhancers required for the retrotransposon transcription^8^. While most hERV loci are inactive due to mutations accumulated through evolution, a subset, most notably the hERV-K (HML-2) subgroup, retains intact open reading frames in its *gag*, *pro*, *pol*, and *env* genes^10^. The HML-2 *pol* locus encodes the essential enzymatic machineries, including a reverse transcriptase (RT)^11^ and an integrase (IN)^12^, which mediate the conversion of the retrotransposon RNA into double-stranded cDNA and its subsequent integration into the host genome, respectively. As the only lineage to have maintained replicative activity throughout recent human evolution, HML-2 exhibits significant insertional polymorphism across contemporary populations^13^.

HERV integration proceeds through a highly orchestrated sequence of molecular events^14^. First, the integrase cleaves two nucleotides from each 3′ end of the cDNA to generate recessed hydroxyl groups (3′-processing). Next, in coordination with divalent magnesium cations (Mg^2+^), these 3′-OH groups execute concerted nucleophilic attacks on the phosphodiester bonds of chromosomal target DNA (tDNA)^14^. These steps result in the formation of new covalent bonds and the strand transfer of the retrotransposon gene into the host genome^14^. These catalytic events occur within a multimeric nucleoprotein complex termed the intasome (Extended Data Fig. 1a), which assembles on paired LTRs and transitions through distinct conformations: the cleaved synaptic complex (CSC), containing 3′-processed viral ends; the target capture complex (TCC), which engages host DNA; and the strand transfer complex (STC), the final state representing the covalently linked integration product^14^.

Notably, emerging evidence highlights the clinical significance of hERV dysregulation as a driver of human pathologies^15^, including oncogenesis^16,17^, neurodegeneration^18^ and aging-related disorders^19,20^. While typically silenced via promoter hypermethylation and histone modifications, the overexpression of HML-2 transcripts and proteins, such as the accessory proteins Rec and Np9^21,22^, is a hallmark of germ cell tumors^23,24^, melanoma^25,26^, and amyotrophic lateral sclerosis (ALS)^27,28^, and it is thought to disrupt host proteostasis and trigger chronic inflammatory cascades^29,30^.

Given the pathogenic potential of these elements, the structural and mechanistic characterization of hERV-K integration activity is of both evolutionary and biomedical significance. However, despite remarkable advances on the elucidation of intasome structures from exogenous retroviruses, including those of HIV-1^31–33^, HTLV^34,35^, the Prototype Foamy Virus (PFV)^36,37^, the Respiratory Syncytial Virus (RSV)^38,39^, the Maedi-Visna Virus (MVV)^40,41^ and the Mouse Mammary Tumour Virus (MMTV)^42,43^, no structural information on human endogenous retroviruses has yet been reported. This represents a major gap in understanding how ancient viral integration machineries have evolved and whether they remain responsive to retroviral inhibitors widely used in antiretroviral therapy.

Here, we present high-resolution cryo-electron microscopy (cryo-EM) structures of the hERV intasome in three distinct states: (i) a complex featuring asymmetric tDNA engagement, (ii) a post-catalytic assembly following strand transfer, and (iii) a dead-end complex bound to an integrase strand transfer inhibitor (INSTI). These structures reveal the molecular architecture of the hERV-K intasome, delineate key protein–DNA and protein–protein interactions driving catalysis, and rationalize how INSTIs restrict hERV catalytic activity. Our findings provide a molecular basis for understanding endogenous retroviral integration and open new avenues for the therapeutic targeting of retrotransposon reactivation in human disease.

## hERV-K intasome adopts a modular architecture with compositional heterogeneity

To elucidate the structural organization of hERV-K (HML-2) integration machinery, we initially sought to reconstitute a cleaved synaptic complex (CSC) *in vitro.* Using recombinant hERV-K108 integrase (IN, Ext. Data Fig. 1b) and oligonucleotide substrates mimicking the LTR termini, the assembly was performed via differential salt dialysis, adapting protocols established for other retroelements. Fractionation by size-exclusion chromatography (SEC) revealed the formation of higher-order nucleoprotein species, displaying an elution profile distinct from that of free IN or LTR DNA (Extended Data Fig. 1c). To resolve the architecture of the hERV-K intasome, we subjected the SEC-purified complex to single-particle cryo-EM. Data were collected on a Titan Krios microscope equipped with a Gatan K3 direct electron detector. A tilted data collection strategy was employed to mitigate preferential particle orientation (Extended Data Fig. 2a-b), enabling a comprehensive sampling of the hERV-K intasome conformational landscape (Extended Data Fig. 2c-d). Unsupervised 3D classification of the cryo-EM data revealed five distinct states (Fig. 1a), with nominal resolutions ranging from 3.40 to 3.76 Å (Extended Data Fig. 2d, Extended Data Fig. 3a-c). Initial atomic coordinates were placed via rigid-body docking of the LTR molecules and discrete IN domains, followed by subsequent refinement of the atomic model. The hERV-K intasome organizes as a two-fold symmetric nucleoprotein assembly and the solved states highlight a noticeable gradient of structural and compositional plasticity. While the complex core remains rigidly defined, the peripheral domains exhibit varying degrees of occupancy and flexibility (Fig. 1a and Extended Data Fig. 3b and 3e).

**Figure 1.**
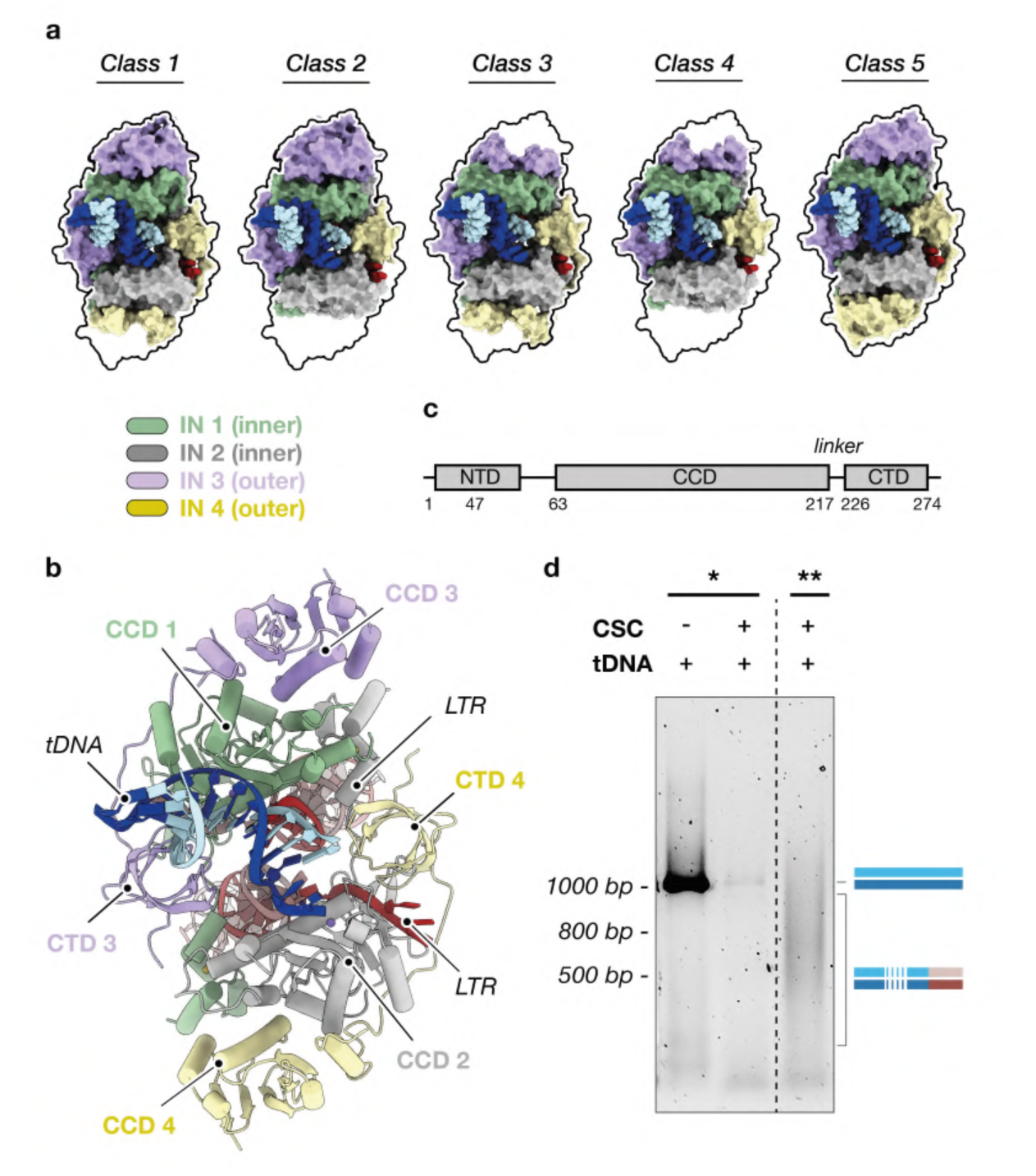
Cryo-EM structures of hERV-K intasome. **a**, Surface representation (top view) of five representative classes of hERV-K intasome complex. Four constitutive integrase (IN) molecules are numbered and colored in green, grey, purple and yellow. The inner or outer positioning of the integrase protomers is specified. Target DNA (tDNA) observed in the DNA binding cleft is colored in blue. Differential domain occupancy is highlighted by a black profile, defined by the outer limits of hERV-K intasome class 5. **b,** Molecular model of hERV-K intasome depicted as cylinders (α-helices) and planks (β-strands), colored as in a. LTR molecules are represented as red ribbons. **c,** Schematic representation of hERV-K integrase domain organization (NTD, N-terminal domain; CCD, catalytic core domain; CTD, C-terminal domain). **d,** Activity assay of hERV-K integration machinery analyzed in an agarose 1% gel. Reconstituted hERV-K intasomes (CSC) were incubated with 1 kb tDNA and the samples were subjected to polymerase chain reaction (PCR) amplification. Oligonucleotides were designed to detect the tDNA template (asterisk) or the retrotransposition products (double asterisk).

Overall, the resulting structures delineate a complex comprising two LTR molecules and up to four hERV-K IN subunits, organized as a dimer-of-dimers architecture (Fig. 1b). The canonical hERV-K IN monomer comprises a three-domain architecture: an N-terminal domain (NTD), a catalytic core domain (CCD), and a C-terminal domain (CTD) (Fig. 1b-c). hERV-K IN lacks the N-terminal extended domain (NED) found in other retroelements such as PFV or the Ty3 retrotransposon, which typically stabilizes the LTR termini^36,37,44^. Likewise, it does not adopt the higher-order complexity seen in octameric or hexadecameric intasomes^39,41,42,45^, representing a more compact integration machinery. The hallmark of the hERV-K intasome is a central catalytic core formed by the close apposition of two inner IN protomers (subunits 1 and 2) with the LTR termini. This core is stabilized by two outer protomers (subunits 3 and 4), which project their CTDs inward to scaffold the synaptic interface (Fig. 1b).

Unexpectedly, all resolved reconstructions displayed additional density within the target DNA (tDNA) binding cleft, which was unambiguously modelled as a short LTR duplex (Fig. 1, Extended Data Fig. 3a). The presence of this molecule highlights the high affinity of the hERV-K intasome for double-stranded DNA. Crucially, this nucleic acid spans only one half of the cleft and engages the complex in a sequence-independent manner, mimicking a substrate and providing further structural stabilization under our reconstitution conditions. Furthermore, cleft occupancy inversely correlates with the flexibility of the IN monomers. In regions where the tDNA molecule is bound, the flanking subunits become more ordered, suggesting that the engagement of the nucleic acid acts as a conformational trigger to rigidify the otherwise mobile peripheral machinery (Fig. 1a; Extended Data Fig. 3e). Ultimately, these structural features define the hERV-K intasome as a highly modular, low-order ensemble. However, the asymmetric binding of tDNA fragments necessitated further functional verification to ensure that the reconstituted machinery remains capable of authentic target DNA engagement and integration activity.

## Functional validation of hERV-K intasome catalytic activity

To validate the catalytic competence of the reconstituted complex, we assessed hERV-K integration activity *in vitro*. It should be emphasized that, since this intasome is assembled using short LTRs rather than a long full-length retrotransposon gene, concerted strand transfer would result in the cleavage of the target DNA scaffold. Therefore, after confirming that the complex binds the nucleic acid through electrophoretic mobility shift assays (EMSAs) (Extended Data Fig. 1d), we evaluated integration activity on a 1 kb linear tDNA using two complementary PCR-based readouts (Extended Data Fig. 1e). First, after incubation with hERV-K intasome, we observed an almost complete depletion of the full-length tDNA PCR signal (Fig. 1d), which suggests that the assembled complex achieves high integration efficiency and leads to tDNA substrate cleavage. Second, to directly identify the integration products we employed primers specific to the LTR and the tDNA 5’ terminus, which would allow us to detect the formation of covalent bonds between the retrotransposon and the substrate (Extended Data Fig. 1e). This approach yielded a heterogeneous distribution of molecules characterized by a diffuse signal rather than discrete bands (Fig. 1d), suggesting that hERV-K integrates stochastically into the host DNA. Therefore, the analysis of PCR-amplified cleavage products confirmed that the reconstituted machinery is fully capable of mediating the strand transfer reaction required for retrotransposition.

To validate this non-specific targeting and to analyze the integration process at the single-molecule level, we subjected the PCR-amplified products to Oxford Nanopore Technologies (ONT) sequencing. After filtering for reads flanked by the tDNA 5’ terminus and the canonical LTR sequence, we identified 171,443 total integration products across 214 unique integration sites. Nanopore sequencing coverage revealed a striking enrichment of strand transfer events at the 5’ end of the target DNA, specifically between nucleotide positions 30 and 50, which featured distinct high-frequency insertion hotspots (Extended Data Fig. 1f). By mapping the expected concerted integration events, separated by a predicted 6-bp target site duplication (TSD), we analyzed the nucleotide sequence flanking the strand transfer sites. Despite the presence of integration hotspots, a preferred consensus motif could not be identified (Extended Data Fig. 1g). This suggests that hERV-K randomly targets the genome in the absence of host regulatory factors, supporting the unspecific binding to the nucleic acid observed in our cryo-EM models. Accordingly, we also detected integration products flanked by dual LTRs. These likely arise from secondary reaction events, or autointegrations, wherein primary strand transfer products serve as subsequent substrates for integration. Collectively, these data underscore the catalytic robustness of the reconstituted machinery and its propensity for promiscuous, non-sequence-specific integration.

## Cryo-EM structure of hERV-K strand transfer complex (STC)

To fully elucidate the structural basis of target DNA engagement and integration, we determined the cryo-EM structure of the hERV-K strand transfer complex (STC). To ensure conformational stability and defined stoichiometry, we reconstituted the STC using branched oligonucleotide substrates that chemically mimic the post-catalytic state, featuring covalent bonds between the LTR 3′ ends and the tDNA. Based on the structural homology between hERV-K and other β-retroviruses such as the Mouse Mammary Tumour Virus (MMTV), we utilized a 6-nucleotide target site duplication to space the integration events. Building on our optimized intasome assembly protocol, we employed a similar reconstitution and purification pipeline to isolate the STC (Extended Data Fig. 1h). Purified complexes were subjected to single-particle cryo-EM, utilizing tilted data collection to overcome potential orientation bias (Extended Data Fig. 4a-c). This strategy allowed us to capture the intasome in a state representing the immediate aftermath of concerted strand transfer, enabling the determination of the hERV-K STC structure at a nominal resolution of 3.02 Å (Fig. 2a, Extended Data Fig. 4d-g).

**Figure 2.**
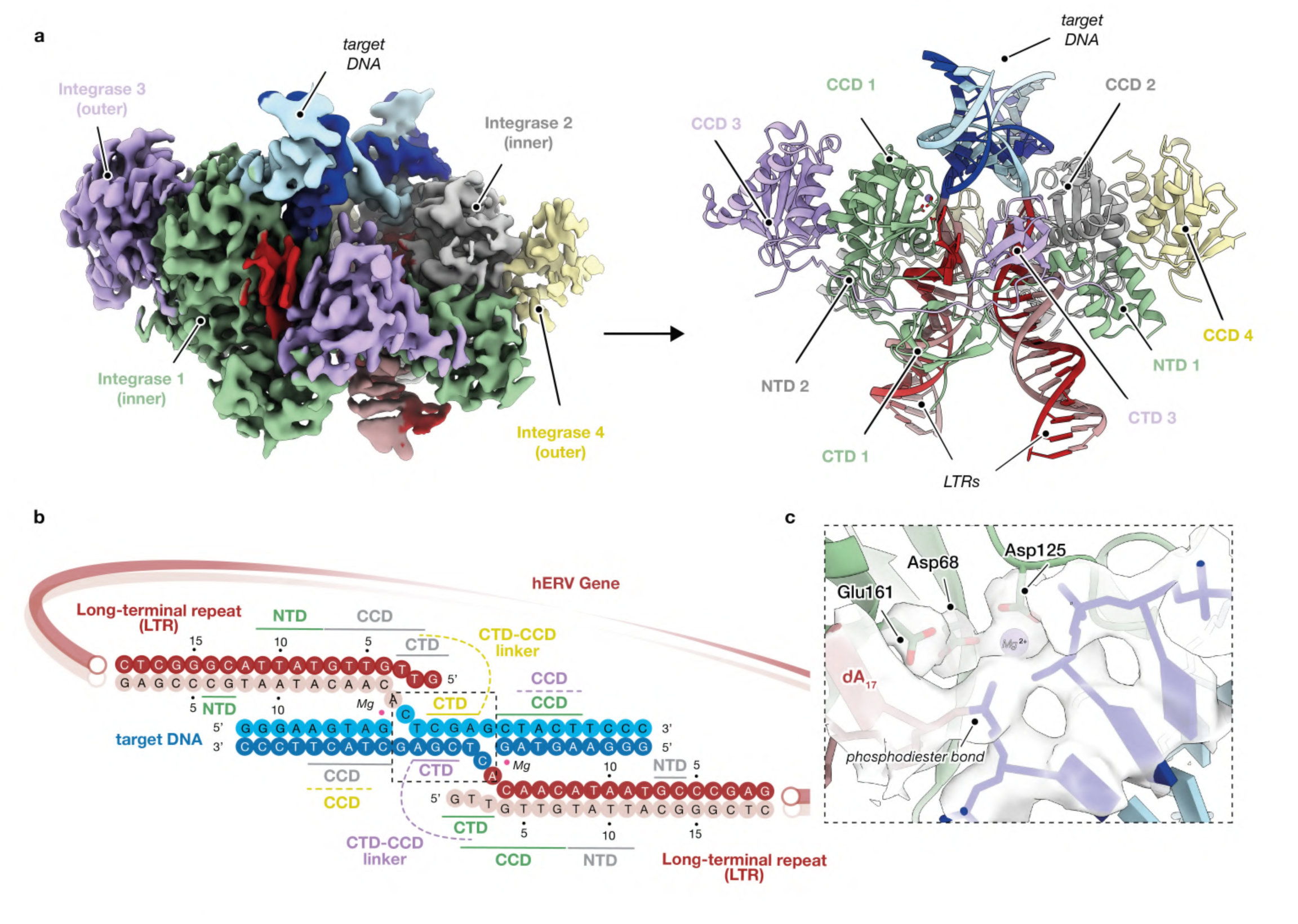
Details of protein-DNA and protein-protein interactions on hERV-K strand transfer complex (STC) **a**, *Left*, Cryo-EM reconstruction of the hERV-K retrotransposon engaged with a target DNA forming a strand transfer complex (STC). hERV-K integrase subunits and LTR molecules are colored as indicated in Fig 1. Target DNA strands are represented in dark and light blue. *Right*, Ribbon representation (side view) of hERV-K STC detailing domain organization and colored as in a. **b,** Schematic representation of STC domain interaction network. DNA nucleotides of the target DNA and LTRs modelled in the hERV-K strand-transfer complex are depicted as solid circles, colored as described before and numbered relative to the nucleic acid 5’ end. Protein-DNA and protein-protein interactions are indicated with solid and dashed lines, respectively. hERV-K integrase subunits are colored as in Fig 1. **c,** Detail of hERV-K intasome active site. Side chains of residues constituting the DDE catalytic triad (Asp68, Asp125 and Glu161) are shown in stick representation. Magnesium ion (Mg^2+^) is represented as a purple sphere. The cryo-EM map surrounding the active site is presented as a grey density, highlighting the quality of the reconstruction on this area.

The hERV-K STC architecture mirrors the primed configuration of the asymmetrically engaged intasome described before (hereafter referred to as hemi-STC), but with full occupancy of the tDNA binding cleft (Fig. 2a). Consistent with the intasome assemblies of PFV^36,37^ and HIV-1^31–33,45,46^, the tDNA is hosted within the central cleft formed by the two-fold symmetric IN dimer-of-dimers (Fig. 2a and Extended Data Fig. 5a-b). At the catalytic or synaptic interface, the tDNA is clamped by the β1–β2 loop of the outer-subunit CTDs (Fig. 2b and Extended Data Fig. 5c), while the peripheral regions are scaffolded by the outer-subunit CCDs (Fig. 2a-b and Extended Data Fig. 5b). This extensive nucleoprotein network rigidifies the strand transfer site, imposing a sharp ∼90° bend in the tDNA trajectory (Extended Data Fig. 5d). Accordingly, CryoSPARC 3D flexible refinement (3D Flex) reveals that while the central integration site is locked in a rigid conformation, the flanking tDNA ends retain significant flexibility (Extended Data Fig. 5e; Supplementary Video 1). This suggests that the hERV-K machinery achieves high-fidelity catalysis by precisely anchoring the target phosphodiester backbone at the point of attack, while allowing the distal DNA to remain dynamic. This structural constraint ensures that the target DNA is presented to the catalytic center with high topological precision, maintaining the specific chemical environment necessary for the strand transfer reaction. The spatial positioning of the dual active sites is physically dictated by the interdomain linkers and the recruitment of the CTDs, which together create a scaffold precisely wide enough to accommodate a 6-bp target site duplication (Fig. 2b, Extended Data Fig. 5d). This integration pattern is a hallmark of β-retroviruses that stands in clear contrast to the 4-bp spacing of spumaviruses (e.g., PFV) or the 5-bp duplication characteristic of lentiviruses (e.g., HIV-1).

The STC structure provides a high-resolution snapshot of the catalytic environment following strand transfer. Within the active sites, the cryo-EM density was sufficiently resolved to define the canonical DDE catalytic triad (Asp68, Asp125, and Glu161) in complex with one Mg^2+^ ion (Fig. 2c). This metal precisely coordinates the newly formed phosphodiester bond, which links the LTR 3′-OH to the target DNA substrate (Fig. 2c). By capturing the post-nucleophilic state, this model confirms that the hERV-K machinery retains the fundamental phosphoryl transfer geometry characteristic of the retroviral integrase superfamily, while utilizing its unique architectural framework to facilitate it.

Despite the high degree of structural stability within the catalytic core, the peripheral regions of the STC reveal a more dynamic nature following the completion of strand transfer. While both pairs of synaptic CTDs remain well-defined within the STC, we observed a noticeable asymmetry in the peripheral domains. Specifically, the outer CCD of subunit 4 exhibits significantly higher flexibility than its subunit 3 counterpart (Fig. 2a and Extended Data Fig. 5f), further supporting the inherent modularity of the hERV-K assembly observed before in the hemi-STC. Consistent with this, the localized disorder correlates with a loss of density for the CTD of inner subunit 2, which lays in close proximity (Extended Data Fig. 5f).

Altogether, these structural insights confirm the hERV-K intasome as a compact archetype of the retroviral integration machinery, where architectural efficiency and catalytic conservation are tightly coupled to ensure the persistence of this ancient retroelement within the human genome.

## Divergent organization of hERV-K and betaretroviral intasomes

Despite the close phylogenetic relationship between hERV-K and exogenous β-retroviruses such as MMTV, our model reveals a striking divergence in intasome organization. While the connectivity of this ensemble is preserved by a complex network of inter-subunit interactions, a defining feature of the hERV-K intasome lies in the spatial arrangement of its outer CTDs (Fig. 3a). In previously characterized octameric MMTV intasomes^42,43^, the synaptic CTDs are donated by peripheral subunits whose associated N-terminal domains (NTDs) and catalytic core domains (CCDs) remain isolated from the nucleoprotein core (Fig. 3a). Accordingly, the CTDs of the outer subunits adopt a scaffolding role, stabilizing the inner CTDs but not participating directly in the enzymatic reaction (Fig. 3a). In contrast, in our hERV-K cryo-EM reconstructions we observe a density bridging the outer subunit CCDs and the synaptic region (Fig. 3b). This suggests that the hERV-K assembly adopts a uniquely compact configuration in which the outer CCD-CTD linker spans the distance to the catalytic interface, thereby positioning the C-terminal domain near the active site (Fig. 3a-b).

**Figure 3.**
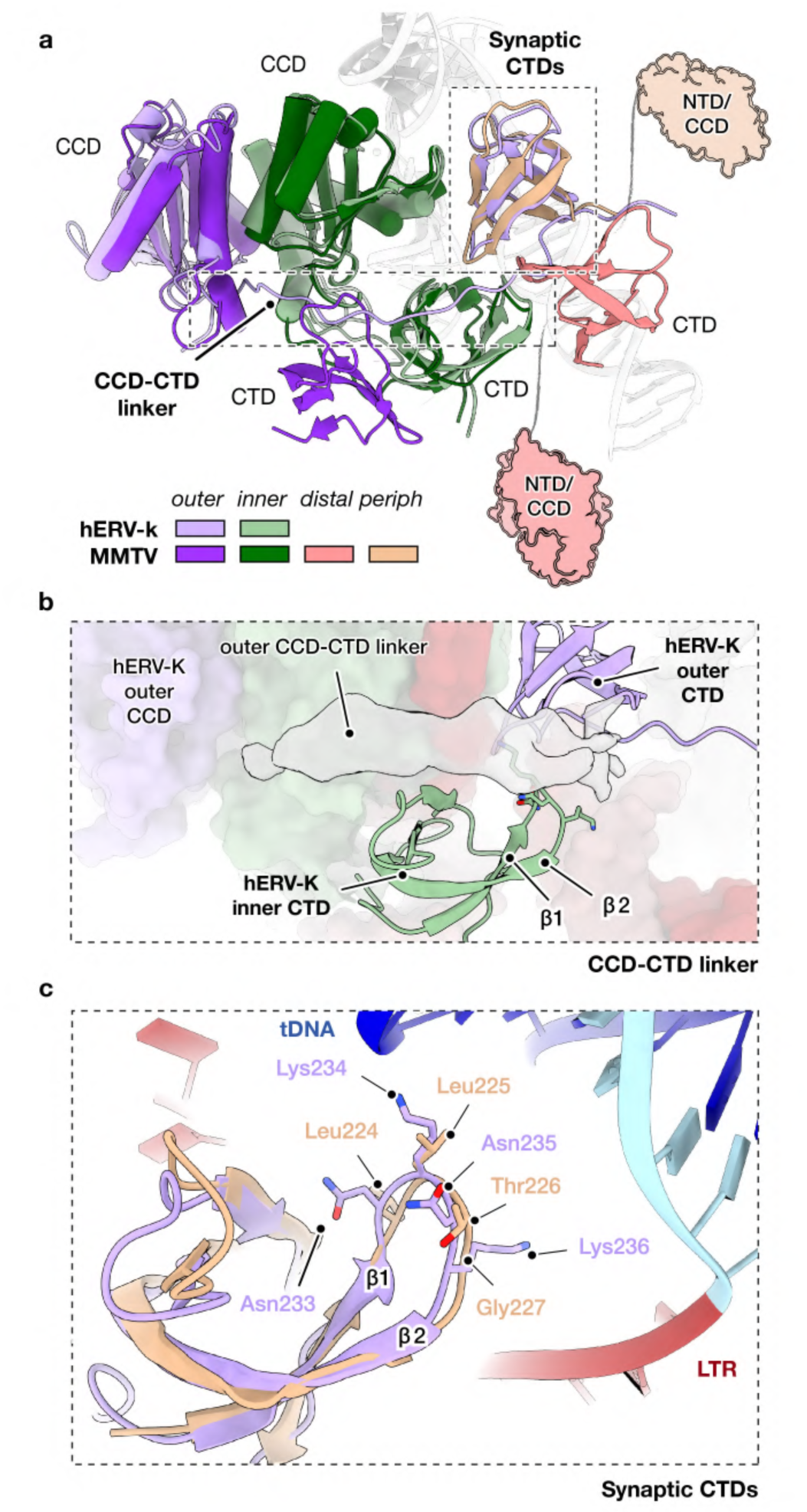
Topological divergence of hERV-K outer CTDs from exogenous retroviral elements. **a**, Human endogenous retrovirus K (hERV-K) shares the general architecture of exogenous betaretroviruses, such as the Mouse Mammary Tumour Virus (MMTV), but adopts a distinct topological domain organization. Structural models are superimposed on the inner catalytic core domain (CCD, green). Color key highlights the relative position of the integrase protomers of each retroelement (inner, outer, distal or peripheral). Position of distal (dark pink) and peripheral (wheat) MMTV domains is represented as cartoon for clarity. **b,** Detail of outer CCD-CTD linker region. Cryo-EM reconstruction (grey density) shows the path adopted by the linker connecting hERV-K outer CCD and CTD, which differs from the orientation observed in MMTV. Molecular models of hERV-K inner and outer CTDs are depicted as purple and green ribbons, respectively. Side-chains of the β1-β2 loop residues are shown. **c,** Ribbon representation of hERV-K (purple) and MMTV (wheat) synaptic CTDs. Side chains of the β1-β2 loop residues are shown as sticks. DNA is represented as ribbon and colored in blue (target DNA) and red (LTRs).

To identify the molecular basis for this architectural divergence, we performed a detailed sequence comparison between the C-terminal domains of hERV-K and MMTV. We identified two critical structural determinants that rationalize the organization of the hERV-K outer subunits. First, in hERV-K, the CTD β1–β2 loop (residues 233-NKNK-236), which is important for stabilization of the newly-formed phosphodiester bond, is composed of large, polar residues, whereas the equivalent MMTV loop (224-LLTG-227) is predominantly hydrophobic (Fig. 3c and Extended Data Fig. 6). In our STC model, these polar hERV-K residues establish an electrostatic surface near the target DNA and the catalytic core, providing a tighter stabilization of the active site that is likely unattainable by the more hydrophobic MMTV homolog (Fig. 3c and Extended Data Fig. 6). Second, we observed a unique structural crosstalk between the CCD–CTD linker of the outer subunit and the inner-subunit CTD. In the hERV-K intasome, the polar β1–β2 loop of the inner CTD directly faces the outer-subunit linker (residues 217-GKKNSPHEGK-226) (Fig. 3b). This linker is not only slightly longer but markedly more polar than the proline-rich MMTV linker (210-PISADPKP-217) (Extended Data Fig. 6). We hypothesize that this highly charged interface acts as a molecular anchor, tethering the outer CCD–CTD linker to the protein core and precisely positioning the outer CTD within the synaptic cleft (Fig. 2a and Fig. 3c). In contrast, the hydrophobic nature of these regions in MMTV likely precludes such anchoring, causing the outer CTD to adopt a non-synaptic position. Consequently, whereas MMTV must resort to an octameric assembly recruiting additional subunits to provide the necessary synaptic CTDs, hERV-K utilizes its specialized sequence motifs to achieve a structurally “compact” tetrameric state. This architectural economy suggests that hERV-K has evolved to maximize synaptic stability within a minimized protein footprint, a trait potentially favored during its long-term co-evolution with the human host.

## A conserved mechanism of pharmacological inhibition in the hERV-K intasome

Given the high degree of conservation within the catalytic core revealed by our structural analysis, we hypothesized that the hERV-K integration machinery would be susceptible to pharmacological inhibition by Integrase Strand Transfer Inhibitors (INSTI). Originally developed against HIV-1, INSTIs function by binding to the intasome active site and sequestering the catalytic Mg^2+^ ions, thereby blocking strand transfer^46–48^. While first– and second-generation INSTIs (e.g., Raltegravir, Dolutegravir) were optimized for lentiviral inhibition, structural studies have demonstrated their broad-spectrum efficacy against distantly related retroelements, including PFV^36^, RSV^39^ and the Simian T-lymphotropic Virus (STLV)^49^.

This cross-species potency has led to the proposal that INSTIs could serve as therapeutic agents for pathologies driven by hERV-K retrotransposition^28^. Although biochemical assays have previously hinted at the sensitivity of hERV-K to these compounds^50^, the molecular basis of this inhibition has remained elusive. Leveraging our robust intasome assembly platform, we sought to define the structural mechanism of hERV-K inhibition, selecting the first-generation INSTI Raltegravir (RAL) as a representative scaffold to probe the drug-target interface.

To functionally validate the susceptibility of hERV-K to pharmacological inhibition, we adapted our *in vitro* strand transfer assay to quantify drug potency via tDNA template depletion. Based on our previous experimental setup, the active intasome mediates concerted integration into the DNA target, effectively cleaving the template and abolishing the specific PCR signal (Extended Data Fig. 1e). Therefore, enzymatic inhibition would be monitored as the recovery of the PCR amplicon. After confirming that the reaction components (RAL and its carrier DMSO) did not interfere with the PCR chemistry itself, we performed a titration of the drug against the assembled intasomes (Extended Data Fig. 1i). We observed a robust, dose-dependent recovery of the tDNA signal, confirming that RAL effectively competes with the catalytic function of the hERV-K intasome and blocks strand transfer under our reconstitution conditions (Extended Data Fig. 1i).

To resolve the molecular determinants of hERV-K inhibition, we performed single-particle cryo-EM analysis of intasomes treated with RAL (Extended Data Fig. 7a-c). Through a 3D classification and refinement pipeline, we determined two high-resolution structures of the drug-bound complex at 2.89 Å and 3.13 Å resolution, respectively (Extended Data Fig. 7d-f). Both reconstructions exhibit exceptional density for the catalytic core, with local resolution reaching 2.5 Å at the drug-target interface (Extended Data Fig. 7g-j).

The first structure, hereafter referred to as CSC/RAL_OPEN_, represents a canonical RAL-bound CSC capturing the drug in its primary inhibitory pose within the synaptic center defined by the two inner integrase protomers (Fig. 4a). In this configuration, the intasome maintains its fundamental tetrameric core, yet the catalytic pocket is significantly reorganized to accommodate the drug. Our high-resolution cryo-EM density unambiguously resolves RAL binding within each active site, where it is positioned to coordinate the two catalytic Mg^2+^ ions (Fig. 4b and Extended Data Fig. 8a). This coordination effectively displaces the catalytic water molecules and precludes the engagement of the tDNA phosphodiester backbone. Specifically, the fluorobenzyl moiety of RAL docks into the hydrophobic pocket functionally reserved for the 3′-terminal adenosine (dA_17_) of the retrotransposon LTR. Consequently, this terminal nucleotide undergoes a pronounced ∼90° rotation out of the active site, a conformational ejection that renders the intasome catalytically inert (Fig. 4b).

**Figure 4.**
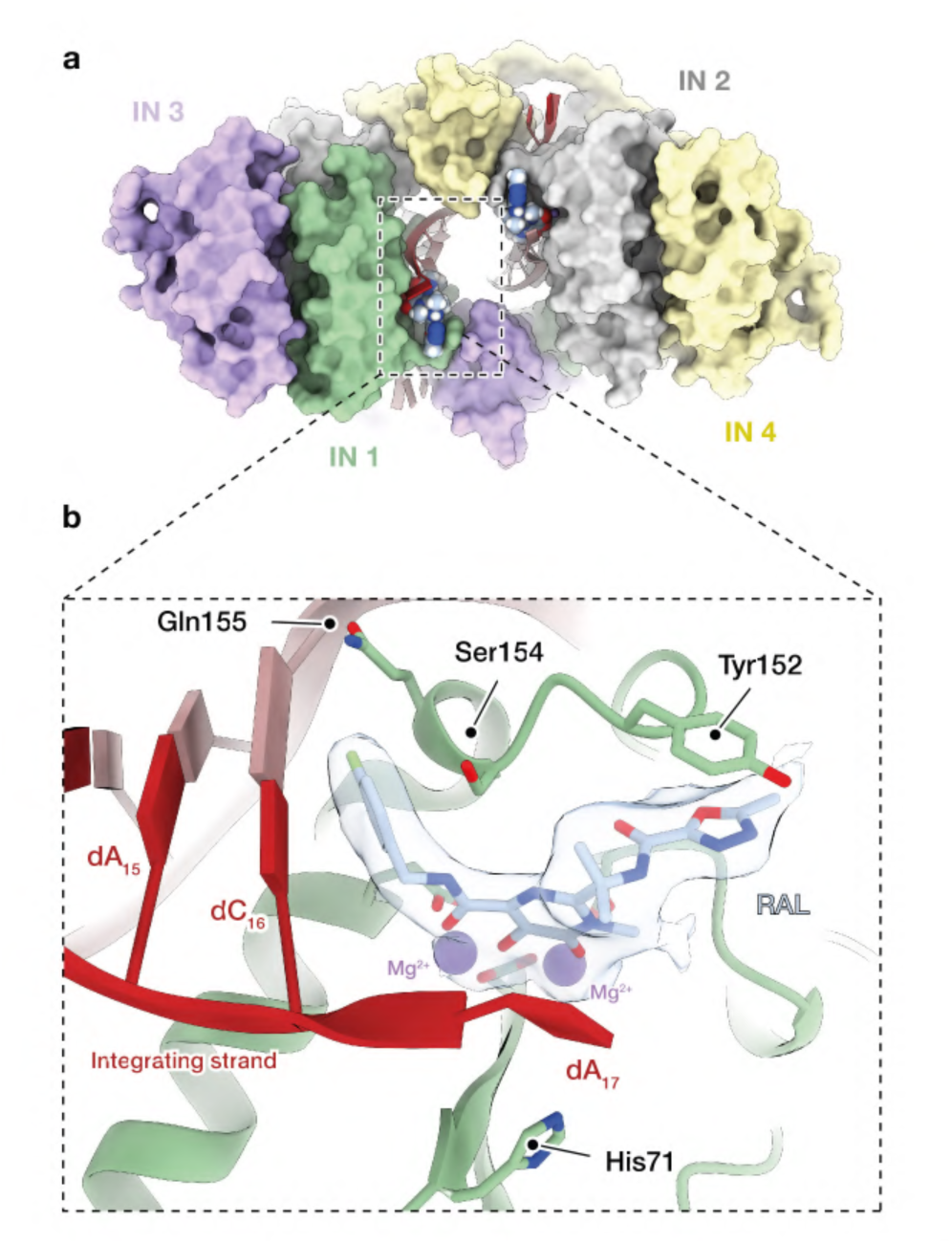
Structural basis of hERV-K intasome inhibition by Raltegravir. **a**, Surface representation (top view) of hERV-K intasome engaged to two Raltegravir (RAL) molecules (depicted as light blue spheres). Integrase protomers are colored as in Fig 1. **b,** Detail of Raltegravir binding pocket. RAL is represented as sticks and colored by atom (oxygen in red, nitrogen in dark blue and fluorine in green). Position of the magnesium atoms is defined as purple spheres. Cryo-EM map around the INSTI drug and catalytic ions is depicted as grey density. Reactive and non-reactive LTR strands are shown as dark and light red cartoons, respectively. Side chains of stabilizing aminoacids are shown as sticks. The fluorobenzyl group of RAL occupies the pocket defined for the 3’-end reactive nucleotide (dA_17_), leading to its displacement and rendering the hERV-K intasome inactive.

The drug-target interface is stabilized by a sophisticated network of π-stacking and polar interactions (Extended Data Fig. 8a). The fluorobenzyl group is sandwiched between the penultimate cytosine (dC_16_) and the side chains of Ser154 and Gln155, while the oxadiazole group establishes a robust π-stacking interaction with the highly conserved Tyr152 (Fig. 4b and Extended Data Fig. 8a). Whereas the homologous position to residue 154 is typically a small hydrophobic proline in other retroviruses (Extended Data Fig. 6 and Extended Data Fig. 8b), the presence of a polar serine in hERV-K does not disrupt the binding pocket; rather, the inhibitor is readily accommodated, suggesting a degree of structural plasticity within the endogenous active site. Finally, the displaced reactive 3′-OH is stabilized in its new position by a stacking interaction between the pyrimidinone core of the drug and residue His71 of the inner IN subunit (Fig. 4b).

Collectively, these structural data confirm that RAL inhibits hERV-K through a mechanism of competitive displacement that is chemically analogous to its action in exogenous retroviruses such as PFV^36^ or STLV^49^ (Extended Data Fig. 8b), notwithstanding the unique architectural framework and specific sequence divergences of the endogenous intasome.

## A drug-stabilized “closed” state reveals an allosteric mechanism of inhibition

Beyond the competitive inhibition mode described above, our cryo-EM analyses uncovered a second, unanticipated structural state of the RAL-bound complex (Extended Data Fig. 7). This alternative architecture, hereafter referred to as the CSC/RAL_CLOSE_ state, is distinguished by a dramatic global reorganization of the synaptic core. Unlike the canonical state, the CSC/RAL_CLOSE_ complex undergoes a significant rigid-body rotation in which one intasome half pivots approximately 50° perpendicular to the CSC symmetry axis (Fig. 5a-b; Supplementary Video 2). This conformational switch results in a severe constriction of the tDNA-binding cleft. Quantification of the solvent-accessible surface (SAS) demonstrates that the cleft narrows from 3101.52 Å² in the CSC/RAL_OPEN_ complex to approximately 611.83 Å² in the closed state (Extended Data Fig. 8c). Functionally, the conformational features defining the CSC/RAL_CLOSE_ state would result in a direct steric clash with the incoming host DNA (Extended Data Fig. 8d), preventing its capture and the subsequent strand transfer reaction. Crucially, this “clamping” mechanism occurs concomitantly with a noticeable increase in the distance between the retrotransposon LTRs (Extended Data Fig. 8e).

**Figure 5.**
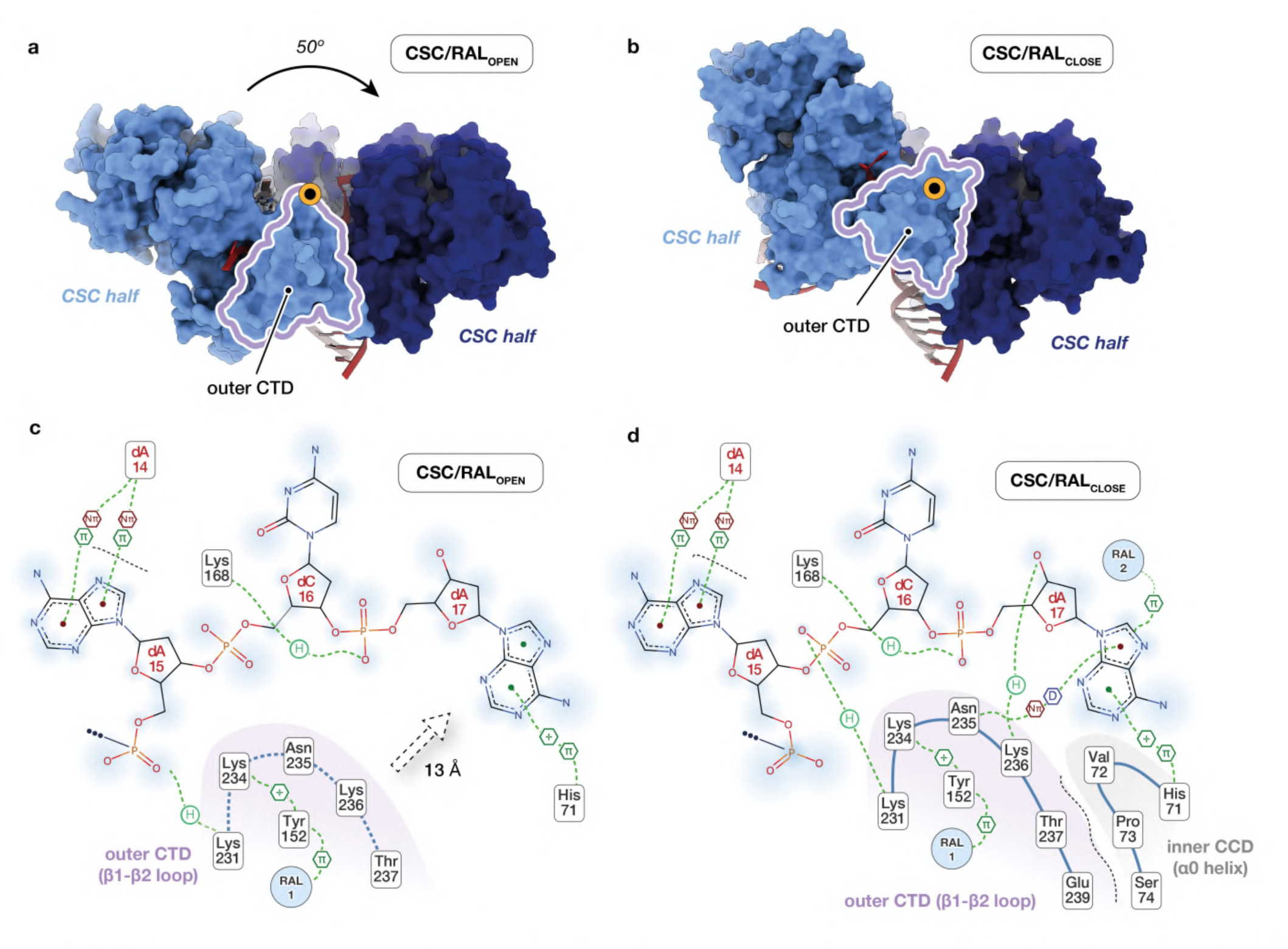
Three-dimensional classification elucidates two high-resolution intermediates of hERV-K inhibition. **a, b**, In the presence of Raltegravir (RAL), hERV-K intasome exists as two distinct conformations, CSC/RAL_OPEN_ (a) and CSC/RAL_CLOSE_ (b), distinguished by a narrowing of the tDNA binding cleft. Driven by this conformational rearrangement, one intasome half (light blue surface) undergoes a 50° rotation relative to the opposing half of the complex (dark blue surface), resulting in the convergence of the outer C-terminal domain (CTD, purple profile) into the opposite active site. Rotation axis is depicted as an orange circle. **c, d,** 2D diagram showing interacting residues, contact network (H, hydrogen bond; D, Donor-π interaction; π, π-π interaction; +, Cation-π interaction) and solvent accessibility (blue shading) of CSC/RAL_OPEN_ (c) and CSC/RAL_CLOSE_ (d) assemblies near hERV-K outer CTD (purple shading). Closure of the tDNA cleft displaces the CTD domain by 13Å, stablishing a novel network of protein-DNA and protein-protein interactions (dashed black line) with the CCD of the opposing inner-subunit (grey shading).

At the local level, the coordination of the Raltegravir scaffold in the CSC/RAL_CLOSE_ state is structurally conserved with the canonical CSC/RAL_OPEN_ complex, preserving the hallmark displacement of the integrating LTR nucleotide (Extended Data Fig. 8f). However, the global architecture is defined by a profound repositioning of the outer-subunit CTD, which undergoes a ∼13 Å translation toward the opposite active site to establish a novel interaction network with the CCD of the opposing inner subunit (Fig. 5c-d). The broader closure is driven by the β1–β2 loop of the outer CTD (Fig. 5c-d, Extended Data Fig. 8f), the same region we previously identified as critical for CCD–CTD linker and active site stabilization, which is notably divergent in the expansive architecture of the MMTV integrase (Extended Data Fig. 6). Thus, the interface between the outer CTD and the LTR is further reconfigured as the β1–β2 loop approximates the integration strand (Fig. 5c-d). Specifically, Lys236, which is solvent-exposed in the open state, is recruited to form a critical hydrogen bond with the 3′-OH (Fig. 5d), and the nucleotide-interaction register of Lys231 is “ratcheted” towards the DNA end, shifting its hydrogen bond from the phosphate of dA_15_ to dC_16_ (Fig. 5c-d).

This closure is further reinforced by a dense network of novel protein-protein interactions between the β1–β2 loop and the α0 helix of the inner CCD. These contacts, not observed in the canonical state, include the engagement of Ser74 (CCD) with Glu239 (CTD), and stabilizing interactions between CTD loop residues (Asn235, Thr237) and core domain residues (Lys168, His71) (Fig. 5d and Extended Data Fig. 8f). Crucially, the α0 helix (residues 72-75), which is critical for CTD stabilization in the CSC/RAL_CLOSE_ conformation (Fig. 5d and Extended Data Fig. 8f), is absent in other retroviruses such as HIV or SIV (Extended Data Fig. 6).

Comparative structural analysis suggests that this docking pocket might represent a regulatory hotspot region. First, in other retroviral CSC models such as those of PFV or STLV, the β1–β2 loop is significantly elongated (Extended Data Fig. 6), and its position correlates within this exact cavity (Extended Data Fig. 8g). Second, re-examination of the hemi-STC cryo-EM reconstructions revealed the presence of a previously unassigned density at the same location (Extended Data Fig. 8g). Remarkably, this extra feature is only found on the half of the tDNA-binding cleft lacking the nucleic acid and it is absent in the fully occupied STC and in the CSC/RAL_OPEN_ state, which hints at a stabilizing role for this region. Altogether, these findings suggest that the CSC/RAL_CLOSE_ state may represent a transient pre-targeting intermediate within the native hERV-K reaction trajectory that is stabilized by RAL binding.

Combined with the observed constriction of the substrate channel, these data indicate that Raltegravir exerts a dual inhibitory mechanism: competitive displacement at the active site and allosteric “locking” of the intasome into a conformation physically incompetent for host target DNA binding and strand transfer.

## Discussion

Although millions of years of co-evolution have rendered most human endogenous retroviruses inactive, their enzymatic competence persists, demonstrated by population polymorphisms and overexpressed hERV proteins in certain pathologies. Our cryo-EM reconstructions reveal the molecular mechanisms of hERV-K integration and rationalize its inhibition by common antiretroviral drugs. In agreement with previous biochemical data, the hERV-K intasome adopts an oligomeric architecture around the long terminal repeats (LTR) of the retrotransposon gene. Its enzymatic core retains fundamental retroelement domain organization, including the conserved catalytic triad and Mg²⁺ coordination. Furthermore, our results confirm the perfect accommodation of a 6-nt TSD, previously only suggested by biochemical assays.

Additionally, our pharmacological analysis confirms hERV-K is susceptible to first-generation INSTIs like Raltegravir (RAL). The canonical binding mode shows that, despite reduced sequence homology, the fundamental inhibition mechanism remains unaltered. Crucially, even with partial tDNA binding cleft occupation, adding RAL to the pre-assembled complex effectively displaces the nucleic acid. This highlights the potency of antiretrovirals to displace pre-engaged tDNA prior to the concerted integration reaction.

Despite these similarities, our results suggest that hERV-K intasome proceeds through a unique assembly and target-recognition mechanism (Extended Data Fig. 9). While other intasomes form large multimeric assemblies (octamers or hexadecamers), our reconstitutions yielded only tetrameric complexes. We hypothesize this lower-order ensemble stems from the unique topology of the outer subunits and synaptic region. Unlike the homologous MMTV intasome, which requires additional monomers to stabilize the integration site, hERV-K performs this function through the C-terminal domain (CTD) of its outer subunits. This compact architecture is unprecedented among retroelements.

Secondly, identifying two distinct drug-bound states suggests the hERV-K CSC might adopt an inactive “pre-engagement” state before tDNA targeting. Initially, the machinery assembles around the hERV-K gene in a closed conformation: the tDNA-binding cleft is constricted while the LTR-binding region is widened to capture incoming LTRs (Extended Data Fig. 9). A shorter electrostatic β1-β2 loop and an α0 helix increase residency time in this “closed” state, allowing us to capture this fleeting conformation. RAL acts as a molecular “wedge”, further trapping this transient state. Following IN engagement with retrotransposon termini, a gating mechanism likely drives the transition to an open conformation: closing the LTR cleft, stabilizing the DNA ends, and opening the tDNA cleft for target recognition (Extended Data Fig. 9). In contrast, in other retroelements, the presence of elongated (e.g., STLV, PFV) or hydrophobic (e.g., MMTV) β1-β2 loops and the lack of an α0 helix (e.g., HIV) render the closed complex highly transient, displacing equilibrium toward an open, tDNA-engagement-prone state. Crucially, the lack of inhibitor-bound structures for related β-retroviruses like MMTV suggests this fleeting state may have remained undiscovered due to the absence of RAL-mediated stabilization.

Structurally, the hERV-K intasome exhibits high peripheral flexibility and, under our reconstitution conditions, reduced solubility compared to other elements. We attribute this instability to its lower-order tetrameric architecture. Consequently, the hERV-K integration machinery is predisposed to rapid target DNA engagement upon assembly, which subsequently stabilizes the complex. This is supported by the compositional heterogeneity, promiscuous binding to “foreign” dsDNA, and increased stability upon full STC tDNA-binding cleft occupancy. Furthermore, the lack of discrete structural sub-classes within our reconstituted STC, alongside minor peripheral subunit differences, underscores the system’s dynamics and propensity for rapid disassembly post-integration.

We propose that this compact organization, high flexibility, and rapid equilibrium between open/closed states result from the ancestral endogenization and co-evolution within the human genome. Physiologically, assembling a lower-order complex requires a minimized protein footprint, favoring host survival. The host-retrotransposon equilibrium also favors less active complexes, as elevated activity risks pathological insertions into essential genes or regulatory elements. Thus, the persistence of the closed state might represent a safer “resting” state for the intasome.

Altogether, our insights provide a structural framework linking hERV-K’s evolutionary adaptation to its unique architecture, strengthening the rationale for using antiretroviral cocktails against hERV-K-associated diseases.

## Methods

### Protein expression and purification

A Histidine14-MBP-SUMO plasmid containing human full-length hERV-K108 integrase was transformed into *E. coli* Rosetta (DE3) pLysS competent cells. Cultures were grown at 37 °C in LB medium supplemented with 50 μg/mL ampicillin until OD_600_ of ∼0.9. Subsequently, cells were incubated at 4 °C for 1 h and underwent overnight induction with 1 mM IPTG and 50 μM ZnCl_2_ at 15 °C.

After cell harvesting, the collected pellet was resuspended in 200 ml lysis buffer consisting of 20 mM HEPES pH 8, 750 mM NaCl, 10 mM MgCl_2_, 10 μM ZnCl_2_, 10 mM Imidazole, 5 mM CHAPS, 10 mM β-mercaptoethanol, 700 ng/ mL Pepstatin A, 50 μg/ml PMSF, 500 ng/mL Leupeptin hemisulphate, 250 μg/mL Benzamidine, supplemented with two protease inhibitor tablets (Thermo Fisher Sci.) and 0.1 mg/ml DNAse.

Then, cells were sonicated (13 cycles, 20 s ON, 59 s OFF, 10 microns amplitude) and the lysate was fractionated via centrifugation at 40,000 x g for 50 min at 4 °C. Afterwards, the soluble fraction was loaded into a HisTrap HP 5 mL column (Cytiva) equilibrated with lysis buffer. After column washing, the protein was eluted in 20 mM HEPES 8, 750 mM NaCl, 10 mM MgCl_2_, 10 μM ZnCl_2_, 300 mM Imidazole, 5 mM CHAPS, and 10 mM β-mercaptoethanol. Fractions corresponding to the hERV-K integrase elution peak were pooled and incubated with SENP1 protease in a 1:60 ratio (w/w) for 20 minutes at 4 °C, to remove the His-MBP-SUMO tag.

The cleaved sample was then diluted with buffer HepA (20 mM HEPES pH 8, 50 μM Na_2_EDTA, 5 mM CHAPS, 10 % Glycerol, 2 mM DTT) until a NaCl concentration of 350 mM. This sample was loaded into a HiTrap Heparin HP 5 mL column (Cytiva), equilibrated with a matching salt concentration. After washing, sample was eluted with a isocratic gradient from 400 mM to 1 M NaCl.

Fractions corresponding to the elution peak were assessed by SDS-PAGE. Those containing pure integrase were pooled, supplemented with arginine and concentrated to ∼4 mg/mL, flash-frozen in liquid nitrogen and stored at –80 °C. The final yield was ∼3 mg protein per culture liter.

### hERV-K intasome Complex Formation

The hERV-K intasome assembly was performed employing a 5’ overhang dsDNA consisting of a 17 bp-long reactive strand (5’– GAGCCCGTAATACAACA-3’; IDT) and a 19 bp-long non-reactive strand (5’– GGTGTTGTATTACGGGCTC-3’; IDT). To obtain double-stranded LTR DNA, both oligonucleotides were heated at 95 °C for 5 min and subsequently cooled down to 12 °C at a –1 °C/min rate. The final dsDNA concentration was 150 μM.

For CSC assembly, 3 mg integrase were mixed with the annealed LTRs in a 1:1 molar ratio at a final concentration of 50 μM. The solution was adjusted to a final concentration of 600 mM NaCl, 10 mM MgCl_2_, 10 μM ZnCl_2_, 1 mM DTT, supplemented with arginine. Following a 2-hour incubation period on ice at 4 °C, the mixture was dialyzed overnight at 4 °C to progressively reduce its salt concentration to 200 mM. The following day, the NaCl concentration in sample was increased to 500 mM and the sample was subsequently incubated for 1 h on ice. Afterwards, the sample was centrifuged to remove any precipitants, and the supernatant was concentrated to 500 μL. Intasome formation was analyzed by size-exclusion chromatography (SEC) in a Superdex 200 increase 10/300 GL column (GE Healthcare), equilibrated with elution buffer (50 mM Tris pH 7, 500 mM NaCl, 10 mM MgCl_2_, 10 μM ZnCl_2_, 1 mM DTT, supplemented with arginine). Fractions corresponding to the first elution peak were pooled.

### Electrophoretic mobility shift assays (EMSA)

The binding of the reconstituted intasome to the target DNA (tDNA) was assessed via electronic mobility shift assay (EMSA). A 34 bp 5’ FITC-labelled DNA strand (5’– /56-FAM/ATATAAAAATTTAAAACTAAGAGAAAAAATCCAA –3’) and its unlabeled complementary strand (5’– TTGGATTTTTTCTCTTAGTTTTAAATTTTTATAT –3’), were annealed as described above, leading to a final dsDNA concentration of 50 μM. Working tDNA concentration was 25 μM.

Pooled intasome fractions were concentrated to A_280_= 0.73. Then, 32 ul intasome or elution buffer (control) was mixed with 1 μl tDNA and incubated for 2 h at room temperature. Following incubation, samples were subjected to electrophoresis at 100 V for 1 h, on a 1 % agarose gel prepared in 1 x TBE buffer. Fluorescently labelled DNA was visualized employing iBright 1500 imaging system (Invitrogen).

### hERV-K Integration/Activity Assays

hERV-K integration activity was assessed by incubating preassembled intasome with tDNA and detecting strand transfer products by PCR amplification.

Integration reactions were initiated by incubating 10 ul of CSC solution with 1 ul of a 1 kb target DNA (50 ng/ul) overnight at RT. Control reactions were performed in parallel using the same buffer conditions as the integration reactions but in the absence of intasome.

Following overnight incubation, the sample was subjected to PCR amplification using Phusion High-Fidelity DNA Polymerase kit (Thermo Fisher Scientific). Two primer sets were used: one pair (set *) flanking both ends of the tDNA sequence (Forward: 5’-GGTACCGGCAAAAAACAGAACC-3’ and Reverse: 5’-CCAAGCTTTACATTTTTGCGTGG –3’) and a second pair (set **) complementary to the tDNA 5’terminus (Forward, described above) and to the LTR non-reactive strand (Reverse: 5’-GAGCCCGTAATACAACA –3’) (Extended Data Fig. 1e).

Following PCR amplification, integration products were resolved by 1% agarose gel electrophoresis at 100V for 1 h and detected with SYBR Safe DNA Gel Stain (Thermo Fisher Scientific) in iBright 1500 (Invitrogen). 100 bp DNA ladder (N3231S, New England Biolabs) was used to identify the size of the resulting DNA products.

### Nanopore Sequencing

To characterize the integration at a molecular level, the PCR products amplified with the second primer set (e.g., tDNA/LTR) were purified using the NucleoSpin Gel & PCR Clean-Up Kit (Macherey-Nagel) and analyzed using Nanopore sequencing.

Libraries were prepared using Ligation Sequencing Kit V14 (SQK-LSK114, ONT) according to the ONT Kit V14 protocol (Document ID: GDE_9161_v114_revAC_24Sep2025, Oxford nanopore) following the manufacturer’s instructions. Briefly, PCR products were subjected to end repair and dA tailing, followed by ligation of sequencing adapters using the NEBNext Ultra II End Repair/dA Tailing Module and Quick T4 DNA Ligase (NEB). Adapter ligated DNA was purified with AMPure XP beads, and the final library was eluted in ONT Elution Buffer (EB; SQK LSK114, ONT).

The prepared library (350ng) was mixed with ONT Loading Beads and Sequencing Buffer according to the SQK LSK114 protocol, and the final mixture was gently loaded onto a primed MinION Flow Cell R10 (FLO-MIN114, ONT).

Sequencing runs were performed in MinKNOW (v24.11.10–22.08.4) using a 16-hour run script with the sequencing kit set to SQK LSK114. Reads were then base called using Guppy (v6.5.7) dna_r10_450bps_fast configuration, and the resulting FASTQ files were subsequently aligned to the reference genome as described below. Upon completion, the flow cell was washed using the Flow Cell Wash Kit (EXP WSH004, ONT) following the manufacturer’s instructions.

### Nanopore Data Analysis

Initial quality control of the high-throughput sequencing reads was performed using Falco, as integrated into the Galaxy v25.1.1.dev0 scientific workflow^51^. This analysis revealed that the MinKNOW sequencing generated a total of 3,756,793 reads with an average length of ∼350 bp.

Following this initial assessment, a custom Python 3 script utilizing the Biopython library^52^ was developed to identify and extract reads representing hERV-K integration events into the DNA substrate. The script computationally generated a library of potential junction sequences by appending the transposon-specific motif (5′-TGTTGTATTA-3′) to the 1 kb template sequence at single-nucleotide increments, starting from a minimum fragment length of 30 bp. Experimental FASTQ files were then screened for exact string matches to these generated sequences or their reverse complements, identifying a total of 94,901 integration-containing reads.

To account for the inherent error rate of Nanopore sequencing, which typically ranges from 2% to 5%, we implemented a fuzzy string-matching algorithm for the identification of integration-site junctions. Given that a standard 40-bp junction (comprising a 30-bp reference prefix and the 10-bp transposon motif) is statistically expected to contain 1–2 sequencing errors, reliance on exact string matching would lead to the erroneous exclusion of a significant proportion of valid reads. By allowing for up to two nucleotide substitutions, we observed a substantial increase in data recovery, with junction detection rising by 81% compared to exact matching (171,443 hits vs. 94,901 hits). To handle the computational load of fuzzy matching across approximately 3.7 million reads, the extraction pipeline was parallelized across 48 CPU cores and executed on a high-performance computing node with 512 GB of RAM, significantly reducing processing time and ensuring a comprehensive search of the integration landscape.

To precisely map the integration sites and analyze the flanking sequence context, a second computational pipeline was employed. First, reads captured on the reverse strand were computationally reverse-complemented to normalize all fragments to the forward reference orientation. Next, the 3′-transposon motif (5′-TGTTGTATTA-3′) was trimmed, and the remaining sequences were locally aligned to the 1 kb DNA template using the Biopython PairwiseAligner (scoring parameters: match = 2, mismatch = –1, gap open = –5, gap extend = –0.5). To reconstruct the sequence context of each integration event while eliminating sequencing noise, the mapped coordinates were used to extract the corresponding 36-bp window directly from the reference template sequence. This window centered the 6-nucleotide tandem site duplication (TSD) between 15 bp of flanking target DNA on each side. To preserve FASTQ formatting for downstream pipelines, these reference-derived target fragments were generated with a uniform, synthetic Phred quality score of 40. These filtered fragments were pooled into a consolidated dataset for downstream characterization of the integration preferences.

Bowtie2 software package^53^, as integrated into the Galaxy scientific workflow, was employed for sequence alignment and analysis, and Unipro UGENE platform^54^ was used for results interpretation

### hERV-K strand transfer complex formation

HERV-K strand transfer complex (STC) was assembled by oligomerization of integrase on a DNA substrate mimicking the strand transfer product. The DNA substrate was generated by annealing three single-stranded DNA oligonucleotides (IDT): (i) a strand corresponding to the 17 bp reactive LTR strand covalently joined to a target DNA sequence, (ii) the complementary non-reactive LTR strand, and (iii) an oligonucleotide corresponding to the tDNA sequence at the 5′ end of the strand transfer junction (Fig. 2b). These oligonucleotides were resuspended as previously described.

STC assembly and isolation were performed following the same methodology as described before.

### hERV-K Activity Inhibition via INSTI drug treatment

hERV-K integration inhibition was assessed by preincubation of the assembled intasome with the integrase strand transfer inhibitor (INSTI) drug Raltegravir (RAL) prior to tDNA addition, followed by PCR-based detection of integration products.

Pooled intasome fractions were diluted to A_280_ = 0.4 in 50 mM Tris-HCl pH 7.0, 5 mM MgCl₂, 1 mM DTT, 200 mM NaCl, supplemented with arginine. Raltegravir working stocks were prepared at 1 mM, 100 µM, 10 µM, or 1 µM by serial dilution of the DMSO stock in Milli-Q H₂O. For inhibitor treatment, 1.1 µL of each RAL working stock was added to 10 µL of intasome solution and incubated for 6 h at RT. DMSO-only control reactions were performed in parallel by adding 1 µL of a 1:10 dilution of DMSO in Milli-Q H₂O. Additional control reactions containing identical buffer conditions in the absence of intasome were included. Following inhibitor preincubation, integration reactions were initiated by addition of 1 µL of tDNA (50 ng/µL) and subsequently processed as described for the integration activity assay.

### Cryo-EM Sample Preparation and Data Collection

Cryo-EM samples were prepared on UltrAuFoil® R1.2/1.3 (300-mesh) gold grids (Quantifoil Micro Tools GmbH). Grids were glow discharged for 3 min at 9 mA using an ELMO glow discharge system prior to application of 4 µL sample at A_280_= 0.73 (hemi-STC) or A_280_= 1.50 (STC). For CSC-RAL samples, intasome was diluted to A_280_= 1.43 in 50 mM Tris-HCl pH 7, 5 mM MgCl_2_ 1 mM DTT, 101.1 mM NaCl, supplemented with arginine and incubated overnight at RT with 32.7 μM Raltegravir. Grids were blotted at 4 °C, 100 % humidity (blot force: 2, blot time: 5.5 s) and plunged frozen into liquid ethane using a Vitrobot Mark IV system (Thermo Fisher Sci.).

Data collections were performed at Instituto Biofisika (Leioa, Spain) on a Titan Krios G4 transmission electron microscope (Thermo Fisher Sci.), equipped with a Gatan Continuum K3 detector (AMETEK, Inc., Pennsylvania, USA), at 300 KeV using the software EPU2. Separate datasets were collected with un-tilted and 30°-tilted parameters to overcome the existence of preferential orientation, which was observed through the presence of anisotropic features in the initial steps of data processing. Collection parameters for each of the datasets have been summarized in Extended Data Table 1.

### Cryo-EM image processing

Frame alignment, anisotropic motion estimation and dose-weighting were performed on-the-fly during data collection using the Patch Motion Correction tool integrated in the CryoSPARC Live suite^55^. Automated Patch CTF estimation was used to estimate the contrast transfer function parameters. Unless stated otherwise, all the subsequent steps of particle picking, extraction, 2D classification, ab-initio reconstruction and map refinements were performed on CryoSPARC v4.7.1^55^.

For the hERV intasome complex dataset, particles were template-picked from the un-tilted and 30°-tilted datasets (Extended Data Fig. 2a-b). This particle set was subjected to multiple steps of 2D classification that provided highly detailed class averages, which accounted for a subset of 651,485 particles (Extended Data Fig. 2c). Next, we performed and *ab-initio* reconstruction at an initial resolution of 35 Å, followed by a Heterogenous refinement step, which resulted in five distinct classes (Extended Data Fig. 2d). Class #4 (233,526 particles) showed defined density features corresponding to an intasome assembly. After further cleaning of the particle set through two-dimensional classification, a Non-Uniform (NU) refinement was performed, which provided a map at 3.23 Å-resolution, according to the gold-standard FSC cut-off criterion. A subsequent heterogeneous refinement step was applied to further eliminate broken or mis-assigned particles, resulting in a final dataset of 187,750 particles that rendered a 3.33 Å-resolution reconstruction after NU-refinement. To prevent misinterpretation arising from the use of C2 symmetry, refinements were performed without imposing any symmetry constraints. Analysis of this cryo-EM map revealed less-defined areas within specific regions of the complex, indicating lower occupancy and/or higher flexibility of the peripheral domains. To address this point, we exported the particle dataset into Relion 5.0 and performed three-dimensional classification without alignment^56^, which rendered five different reconstructions characterized by distinct subunit composition and domain features. Each of the obtained classes was re-imported into CryoSPARC and subjected to local refinement, using the previously defined angular assignments. The resolution of the generated reconstructions spanned from 3.40 to 3.76 Å, estimated at the gold-standard 0.143 cut-off criterion (Extended Data Fig. 3a-c). Local resolution estimation was performed from the output of the refinement jobs (Extended Data Fig. 3b).

For the hERV-K STC dataset, merging of the un-tilted and 30°-tilted micrographs was followed by recursive 2D classification of the template-picked particles, resulting in a 1,596,132-particle set (Extended Data Fig. 4). Subsequent *ab-initio* reconstruction and heterogeneous refinement yielded eight distinct classes, effectively separating low-resolution particles and anisotropic reconstructions from those representing high-quality STC complexes. Among the latter, Class #3 (434,391 particles) was further subjected to non-uniform and local refinement processes, which rendered a 3.02 Å-resolution reconstruction according to the gold-standard cut-off (Extended Data Fig. 4d-f). Local resolution estimation was performed from the output of the refinement job (Extended Data Fig. 4g).

For the RAL-treated intasomes, we followed a similar data-processing strategy. Briefly, after merging un-tilted and tilted micrographs, template-picking and 2D classification yielded a final dataset consisting of 617,079 particles (Extended Data Fig. 7c). *Ab initio* and heterogeneous refinement steps resulted in eight different three-dimensional reconstructions, two of which (Class #0, 181,815 particles; Class #1, 104,815 particles) corresponded to CSC/RAL complexes in distinct conformational states (Extended Data Fig. 7d). Next, non-uniform and local refinement of each of these classes, followed by local resolution estimation, yielded canonical CSC/RAL_OPEN_ and non-canonical CSC/RAL_CLOSE_ reconstructions at nominal resolutions of 2.89 Å and 3.13 Å, respectively, according to the gold-standard 0.143-FSC threshold (Extended Data Fig. 7e-h).

### Cryo-EM model building and refinement

Initial structural models of hERV-K integrase were predicted using the AlphaFold 3 server^57^. To generate a starting model for the hERV-K intasome, these predictions were superimposed onto the octameric cleaved synaptic complex (CSC) structure of the Mouse Mammary Tumour Virus (MMTV) (PDB code: 3JCA)^42^, with which it shares high structural homology. The DNA coordinates from the MMTV template were retained, and the sequence was mutated to match the LTR used in our experimental reconstructions. The resulting *in silico* assembly was divided into its constituent N-terminal (NTD), catalytic core (CCD), and C-terminal (CTD) domains. These individual domains were subsequently rigid-body fitted into the cryo-EM reconstruction of hERV-K CSC using UCSF Chimera (version1.10.1) to establish the initial coordinates and global orientation of the complex^58^. Manual model building was performed in Coot (version 0.9.8.96)^59^, which revealed the absence of the peripheral subunits previously observed in the MMTV intasome. Following manual correction of the coordinates, the model was subjected to multiple rounds of real-space refinement using the Phenix software suite (version 1.21.2-5419)^60^. This iterative process was continued until a complete tetrameric model of the hERV-K intasome was obtained.

At this stage, well-defined unassigned density consistent with a helical structure was still observed near the intasome tDNA-binding cleft. Given the location and morphology of this feature, a double-stranded LTR molecule was rigid-body fitted into the map using UCSF Chimera. Subsequent refinement in Coot and Phenix, as previously stated, facilitated the characterization of an dsDNA-binding event within one side of the cleft. Additionally, an unidentified density was observed bridging the outer integrase subunits and the synaptic CTD region. This feature was attributed to the CCD-CTD linker connecting the two domains. Although the map quality in this region was limited, the corresponding residues were tentatively modelled and manually built in Coot to satisfy the observed density.

Model building of hERV-K STC was performed through the structural superimposition of the MMTV STC model^43^ (PDB code: 7USF) onto the hERV-K intasome core described above. The nucleotide sequence of the target DNA was subsequently mutated to match the experimental sequence, which allowed to stablish the global position and orientation of the nucleic acid within the binding cleft. As described above, manual rebuilding and iterative refinement were performed in Coot and Phenix, to determine the final hERV-K STC molecular model.

Finally, modelling of the RAL-bound structures required two distinct strategies. First, to correctly position the drug within the canonical CSC/RAL_OPEN_ reconstruction, a rigid body fit of a RAL-bound Prototype Foamy Virus (PFV) inner integrase subunit^61^ (PDB code: 3OYA) was performed against the hERV-K STC model described above (with the tDNA molecule removed). This established the initial coordinates of two RAL molecules near the active site, which were further curated through Phenix real space refinement. On the other hand, determination of the CSC/RAL_CLOSE_ atomic coordinates accounted for the significant conformational rearrangement observed in the cryo-EM reconstruction, which precluded a direct model building process. To address these structural differences, the CSC/RAL_OPEN_ model was partitioned into two halves, each of which was rigid-body fitted into the CSC/RAL_CLOSE_ cryo-EM map. This yielded a preliminary CSC/RAL_CLOSE_ model that was further refined in Coot and Phenix to address the specific conformational changes relative to the CSC/RAL_OPEN_ state.

Cryo-EM figures were prepared using UCSF Chimera 1.19^58^ and ChimeraX-1.10.1^62^. Multiple sequence alignments were calculated using the Clustalw Omega server (https://www.ebi.ac.uk/Tools/msa/clustalo/) and figures were prepared in Jalview 2.11.5.1. Structural comparisons were performed in Pymol 3.1.6.1 and UCSF Chimera 1.19^58^. Surface topography and solvent accessibility analysis was performed on the CASTpFold server (https://cfold.bme.uic.edu/castpfold/) using a probe radius of 2.5 Å^63^. 2D interaction diagrams were generated using SAMSON.

## Acknowledgements

This work has been supported by grants PID2021-126263OA-I00 and PID2024-158879NB-I00 to G. A.-P, funded by the Spanish Ministry of Science and Innovation (MICIU/AEI/10.13039/501100011033) and by FEDER, UE.

A. G. A.-P. acknowledges financial support from a Ramón y Cajal program grant RYC2020-029163-I, funded by MICIU/AEI/10.13039/501100011033 and by the European Social Fund (ESF) ‘The ESF invests in your future’. A. B.-M. acknowledges financial support from the Basque Government predoctoral program (PRE_2024_1_0132).

We thank F. Moro for advice during complex reconstitution, A. Jiménez for the computing infrastructure, and all the members of the Abascal-Palacios lab for critically reading the manuscript and for fruitful discussions. We thank A. Gonzalez, I. Sánchez and H. Liang for their support during cryo-EM sample preparation. We thank the Proteomics platform and the Electron Microscopy and Crystallography platform from CIC-bioGUNE for experimental support during the development of this research. We thank the Basque Resource for EM (BREM) supported by the IKUR 2023 strategy of the Department of Science, Universities and Innovation and the Innovation Fund of the Basque Government, with additional support from MCIN (Recovery, Transformation and Resilience Plan) and the Basque Government ‘Biotechnology Complementary Plan Applied to Health’ with funding from European Union NextGenerationEU (PRTR-C17.I1; PRTR-C17.I01.P01.S13; AAAA_ACG_AY_2539/22_05).

## Author contributions

A. B.-M. carried out protein expression and purification, complex reconstitution, activity/inhibition assays, EM specimen preparation and EM data collection. S. F. carried out construct cloning, protein expression and purification and complex reconstitution. M. D.-M. prepared Nanopore libraries, performed integration product sequencing and helped with data analysis. G. A.-P. designed and supervised research, carried out protein expression, purification and complex reconstitution, EM data collection and processing, model building and refinement, analyzed the structural data and wrote the manuscript with contributions from all authors.

## Correspondence

Correspondence and requests for materials should be addressed to Guillermo Abascal-Palacios

## Code availability

The custom Python 3 pipeline used for the extraction, orientation normalization, and reference-correction of hERV-K integration sites is publicly available. The source code will be deposited without restriction in a public GitHub repository and permanently archived on Zenodo upon manuscript acceptance, and is available to reviewers upon request.

## Data Availability

The data that support this study are available from the corresponding authors upon reasonable request. The atomic coordinates and corresponding electron microscopy maps have been deposited in the Protein Data Bank (PDB) and Electron Microscopy Data Bank (EMDB), respectively. The accession codes are: PDB 31BR (pdb_000031BR) and EMD-58258 for hemi-STC class #1; PDB 31BS (pdb_000031BS) and EMD-58259 for hemi-STC class #2; PDB 31BT (pdb_000031BT) and EMD-58260 for hemi-STC class #3; PDB 31BU (pdb_000031BU) and EMD-58261 for hemi-STC class #4; PDB 31BV (pdb_000031BV) and EMD-58262 for hemi-STC class #5; PDB 31BX (pdb_000031BX) and EMD-58263 for STC; PDB 31BY (pdb_000031BY) and EMD-58264 for CSC/RAL_OPEN_; and PDB 31BZ (pdb_000031BZ) and EMD-58265 for CSC/RAL_CLOSE_.

Additional atomic models used in this study correspond to the following RCSB database codes (https://www.rcsb.org/): MMTV cleaved synaptic complex (3JCA), MMTV strand transfer-complex (7USF), PFV/RAL (3OYA), STLV/RAL (7OUG). Source data are provided with this paper.

## Extended Data Figure Legends

**Extended Data Figure 1.**
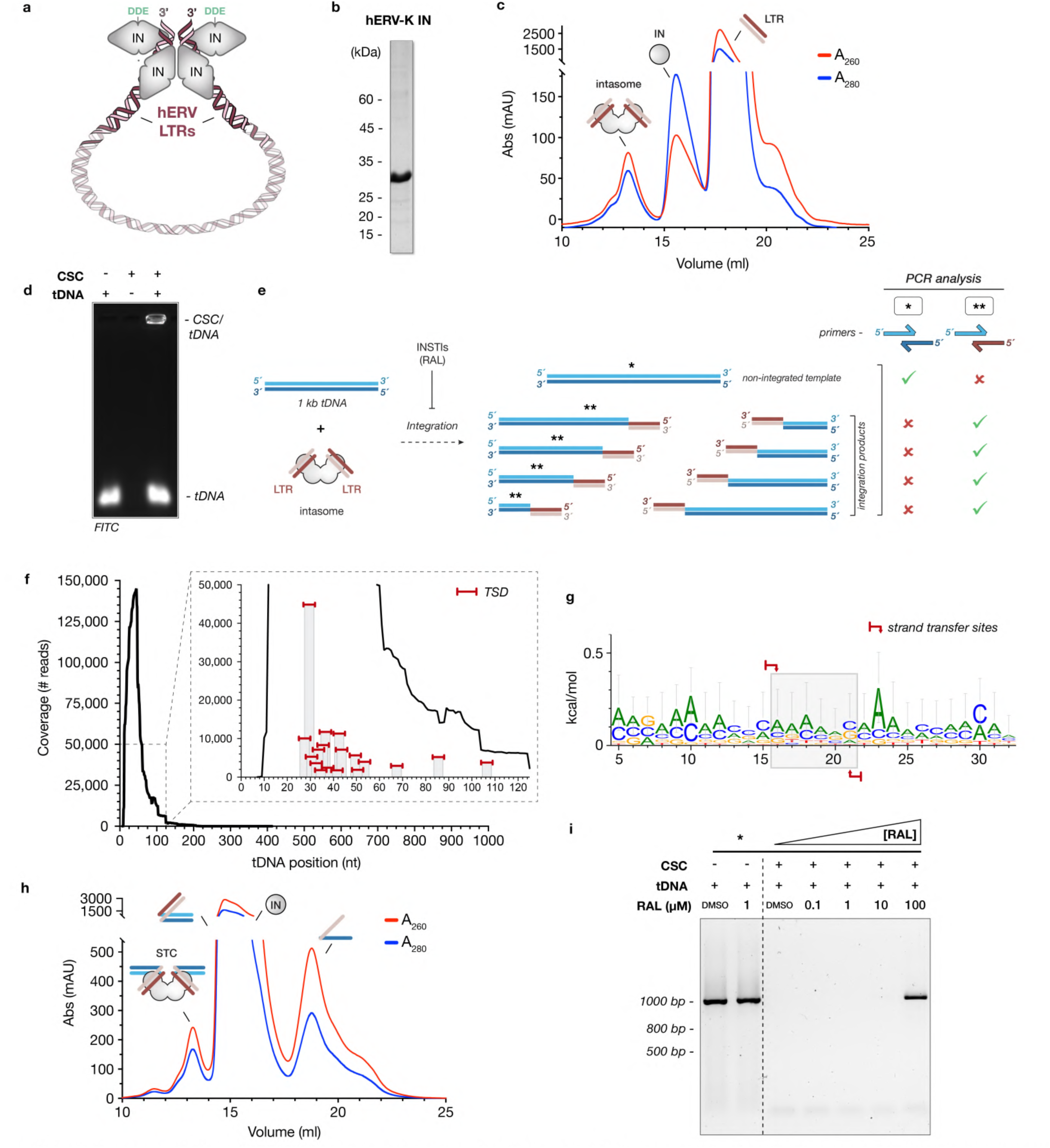
Reconstitution of hERV-K intasome machinery, activity and inhibition assays. **a**, Schematic representation of hERV-K intasome. Integrase subunits are depicted as grey geometrical shapes and retrotransposon gene as red-shaded helices. Long-terminal repeats (LTRs) used on the reconstitution assays are shown in solid colors whereas the absent gene body is depicted as semi-translucent for clarity. Active site is defined by the 3’-hydroxyl group and the catalytic DDE triad (green). **b,** Purification of hERV-K integrase (IN). Representative band from SDS-PAGE analysis is shown. **c,** Size-exclusion chromatography (SEC) of hERV-K intasome. UV absorbance at 260 and 280 nm are indicated as red and blue lines, respectively. **d,** Electrophoretic mobility shift assay (EMSA) demonstrates the ability of the reconstituted intasome to bind to target DNA (tDNA) in-vitro. **e,** Schematic diagram of PCR-based integration assay protocol. Following incubation of the intasome and a 1 kb tDNA, the sample was subjected to two independent PCR reactions to identify non-integrated template (asterisk) or integrations products (double asterisk). **f,** Nanopore sequencing coverage across the 1 kb reference tDNA. The Y-axis indicates the number of independent reads mapping to each nucleotide position. *Inset*, representative hotspots (> 1%) of frequent transposon integration sites. The expected 6-nt tandem duplication sites (TSD) are represented as red bars. **g,** Sequence logo of enriched integration sites. The position of the concerted covalent bonds is depicted as red arrows and delimit the TSD (grey shading). **h,** Size-exclusion chromatography (SEC) of hERV-K strand transfer complex (STC), depicted as in panel c. **i,** PCR-based inhibition assay. Amplification of a 1 kb tDNA (using primer set * from panel e) was employed as a reporter of Raltegravir (RAL) inhibition. Increasing concentrations of RAL were tested.

**Extended Data Figure 2.**
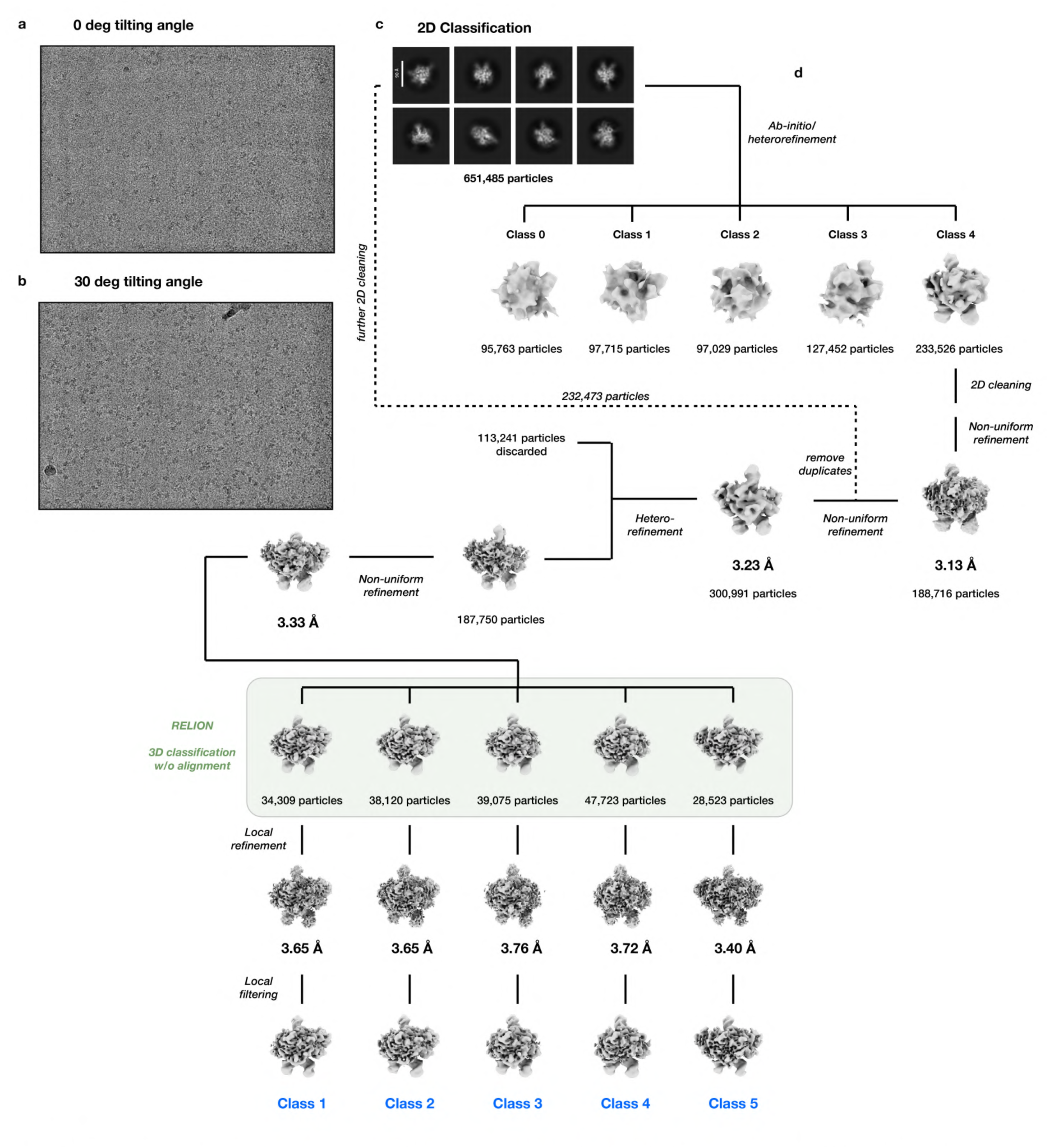
Cryo-EM data processing and resolution estimation of hERV-K intasome complex. **a, b**, Representative raw micrographs of hERV-K intasome collected at 0° tilting angle and 30° tilting angle. **c,** Eight reference-free 2D class averages obtained from 651,485 independent particles. **d,** 3D classification of the joined set of particles from 0° and 30° tilting angles datasets. The particles were subjected to a hierarchical process, encompassing several rounds of classification and refinement in CryoSPARC, as described in the schematic. Particles resulting from non-uniform refinement were exported into Relion (green box) and subjected to further global 3D classification without alignment. The obtained 3D reconstructions and their corresponding particles were re-imported into cryoSPARC and subjected to local refinement and local filtering. The estimated resolution at the gold-standard FSC (FSC= 0.143) and the number of particles contributing to each class are indicated close to the corresponding 3D maps.

**Extended Data Figure 3.**
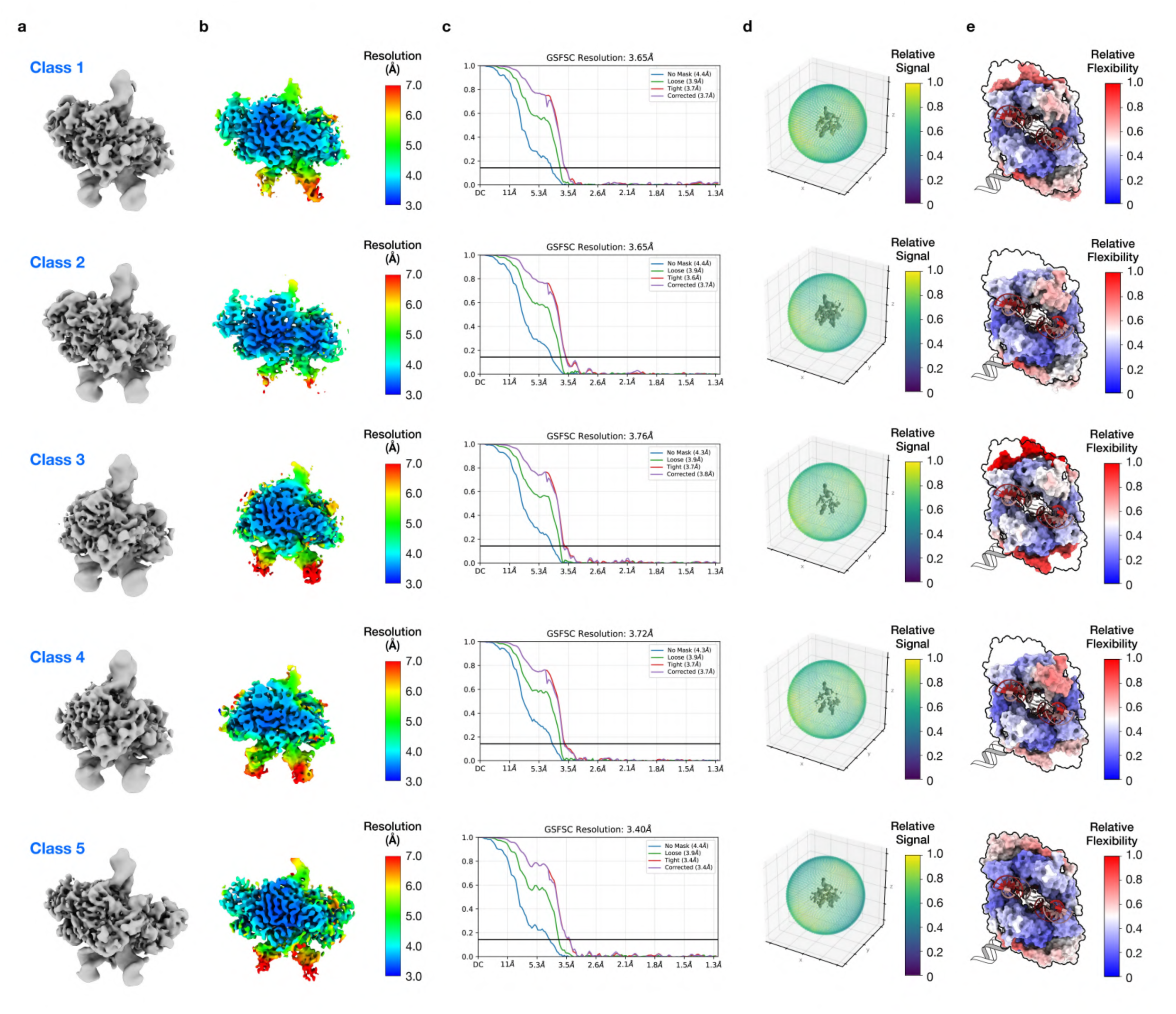
Cryo-EM reconstructions and model fitting of hERV-K intasome complex For each hERV-K intasome class: **a**, Local-filtered 3D reconstruction (lateral view). **b,** Resolution estimation of the cryo-EM map calculated with CryoSPARC. Central slice (*right*) view is shown and colored according to the local resolution, as indicated in the scale bar. **c,** Fourier-shell correlation (FSC) representation of the masked (purple) and unmasked (blue) cryo-EM reconstruction with the estimated resolution at the gold-standard FSC. **d,** Orientation distribution sphere of the particles that contributed to the final cryo-EM reconstruction colored according to the relative signal, as indicated in the scale bar. e, Surface representation (bottom view) of molecular model colored according to its relative flexibility, as indicated in the scale bar. tDNA asymmetric binding position, which is hidden on this orientation, is indicated through a helical representation.

**Extended Data Figure 4.**
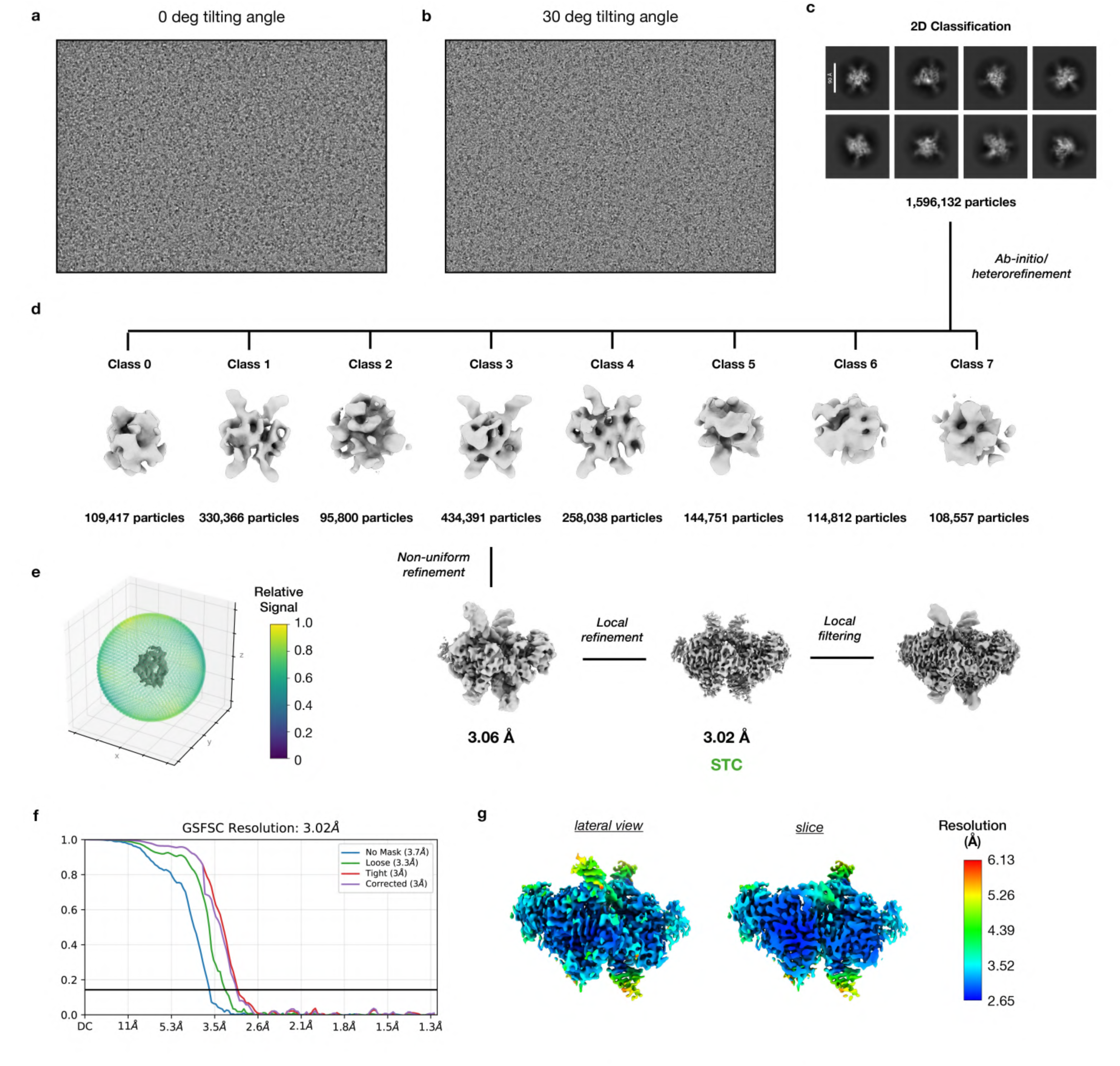
Cryo-EM data processing and resolution estimation of hERV-K strand transfer complex (STC) **a, b**, Representative raw micrographs of hERV-K strand transfer complex (STC) collected at 0° tilting angle and 30° tilting angle. **c,** Eight reference-free 2D class averages obtained from 1,596,132 independent particles. **d,** 3D classification of the joined set of particles from 0° and 30° tilting angles datasets. The particles were subjected to a hierarchical process, encompassing several rounds of classification and refinement in CryoSPARC, as described in the schematic. The estimated resolution at the gold-standard FSC (FSC= 0.143) and the number of particles contributing to each reconstruction are indicated close to the corresponding 3D maps. **e,** Orientation distribution sphere of the particles that contributed to the final cryo-EM reconstruction according to the relative signal, as indicated in the scale bar. **f,** Fourier-shell correlation (FSC) representation of the masked (purple) and unmasked (blue) cryo-EM reconstruction with the estimated resolution at the gold-standard FSC. **g,** Resolution estimation of the cryo-EM map calculated with CryoSPARC. Lateral (*left*) and central slice (*right*) views are shown and colored according to the local resolution, as indicated in the scale bar.

**Extended Data Figure 5.**
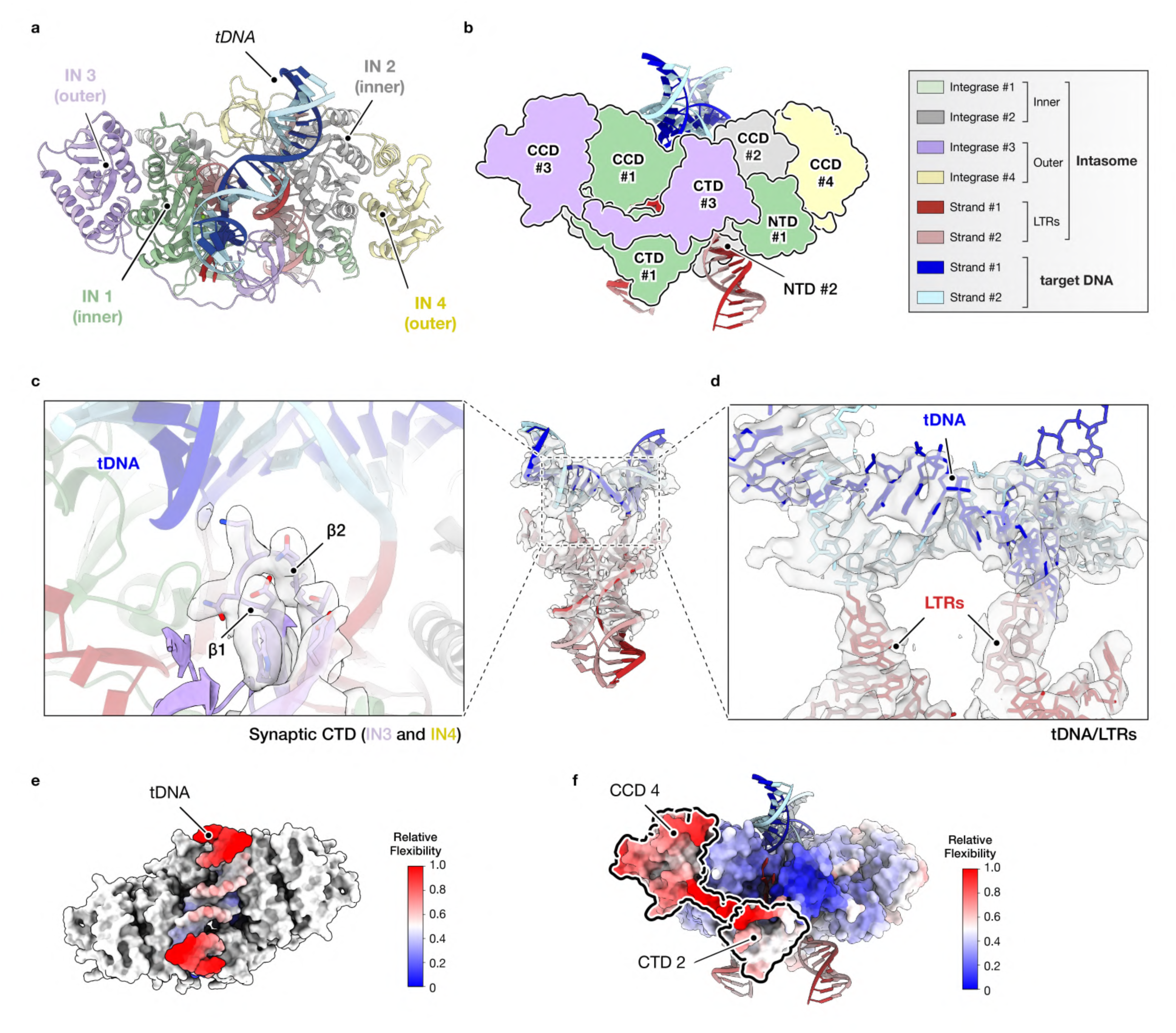
Cryo-EM reconstructions and model fitting of hERV-K strand transfer complex (STC) **a**, Ribbon representation (top view) of hERV-K strand transfer complex (STC) structure model. Subunits are colored and numbered as in Fig. 1: inner integrases 1 and 2 in green and grey, respectively; and outer integrases 3 and 4 in purple and yellow, respectively. **b,** hERV-K retrotransposon STC diagram representing as silhouettes the domain architecture of hERV-K integrase subunits: NTD, N-terminal domain; CCD, catalytic core domain; CTD, C-terminal domain. **c,** Detail of hERV-K synaptic CTD (purple ribbon), highlighting β1-β2 loop engagement at the catalytic interface. Cryo-EM reconstruction is shown in grey. **d,** DNA geometry at STC integration site. LTRs and target DNA are represented as sticks in red and blue shades, respectively. Cryo-EM map is shown in grey. **e,** Target DNA (tDNA) distal region presents higher flexibility. hERV-K intasome molecular model (top view) is depicted as a surface representation, the integrase protomers are colored in grey and the tDNA is colored according to its relative mobility, as indicated in the scale bar. **f,** The flexibility of the outer IN subunits (i.e. CCD 4) directly influences the stability of the inner subunit CTDs (i.e. CTD 2). STC core is colored according to its mobility and representative domains are highlighted with a black silhouette.

**Extended Data Figure 6.**
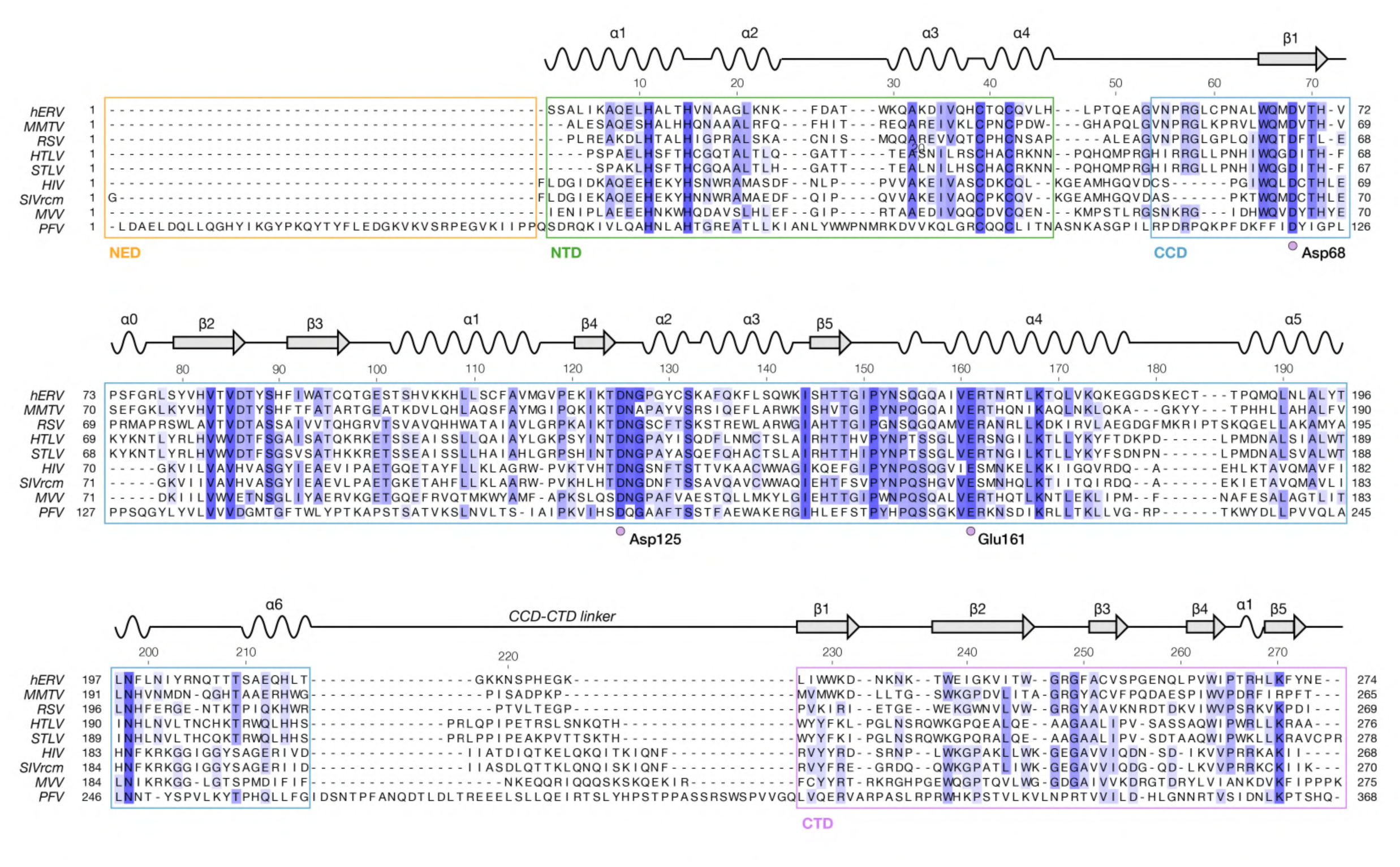
Sequence comparison of hERV-K intasome to exogenous retroviral elements Structure-based sequence alignment of integrase proteins from human endogenous retrovirus K (hERV-K), Mouse Mammary Tumour Virus (MMTV), Respiratory Syncytial Virus (RSV), Human T-lymphotropic virus type 1 (HTLV-1), Simian T-lymphotropic virus (STLV), Human Immunodeficiency Virus (HIV), Simian Immunodeficiency Virus (SIVrcm), Maedi-Visna virus (MVV) and Prototype Foamy Virus (PFV). Secondary structure motifs are represented above each alignment. N-terminal extended domains (NED), N-terminal domains (NTD), catalytic core domains (CCD) and C-terminal domains (CTD) are delimited by orange, green, blue and purple squares, respectively. The position of the DDE catalytic triad is defined by purple circles and numbered according to the hERV-K reference.

**Extended Data Figure 7.**
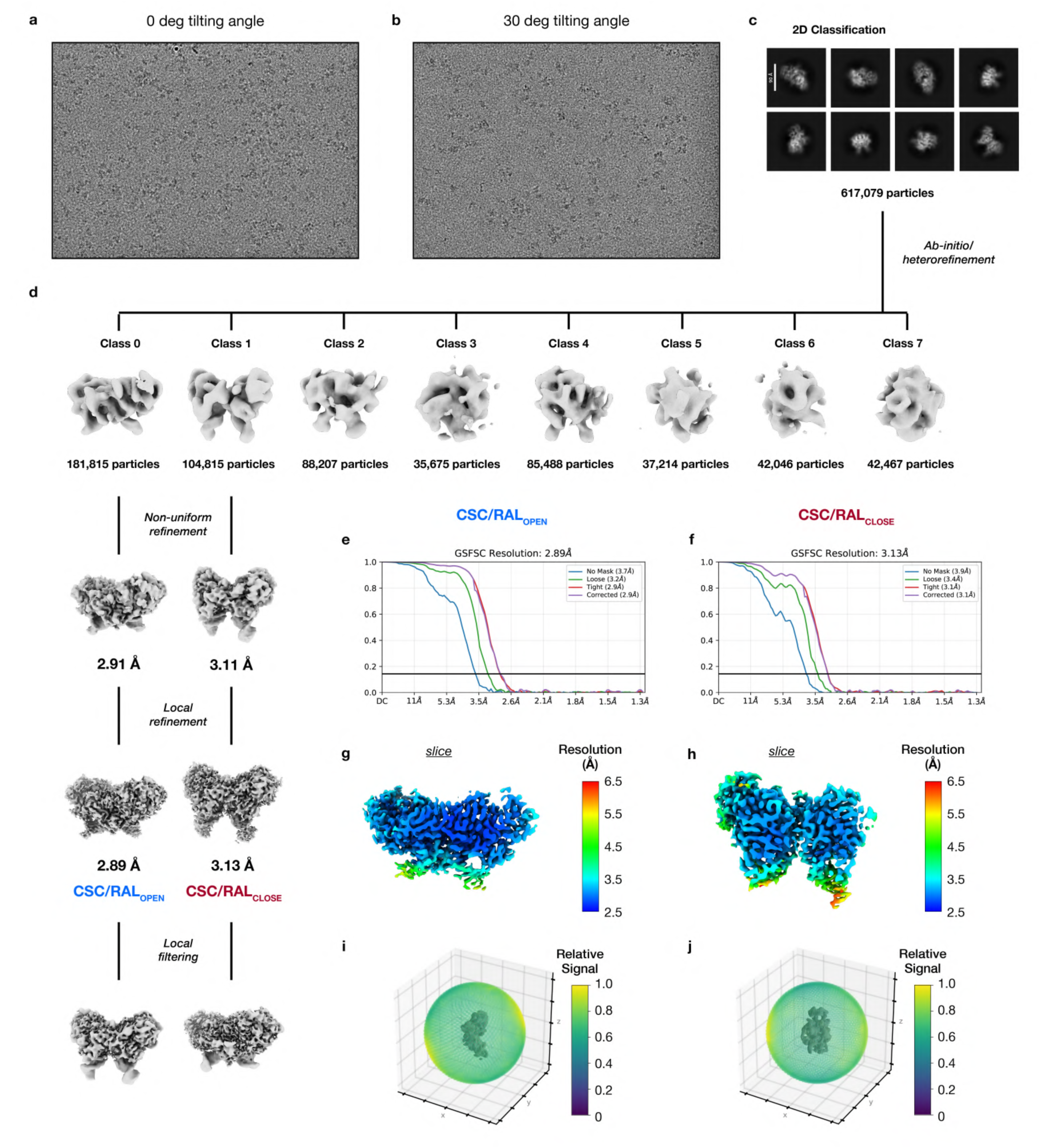
Cryo-EM data processing and resolution estimation of hERV-K intasome bound to Raltegravir (RAL) **a, b,** Representative raw micrographs of hERV-K intasome bound to Raltegravir (RAL) collected at 0° tilting angle and 30° tilting angle. **c,** Eight reference-free 2D class averages obtained from 617,079 independent particles. **d,** 3D classification of the joined set of particles from 0° and 30° tilting angles datasets. The particles were subjected to a hierarchical process, encompassing several rounds of classification and refinement in CryoSPARC, as described in the schematic. The estimated resolution at the gold-standard FSC (FSC= 0.143) and the number of particles contributing to each class are indicated close to the corresponding 3D maps. **e, f,** Fourier-shell correlation (FSC) representation of the masked (purple) and unmasked (blue) cryo-EM reconstruction of CSC/RAL_OPEN_ and CSC/RAL_CLOSE_ with the estimated resolution at the gold-standard FSC. **g, h,** Resolution estimation of CSC/RAL_OPEN_ and CSC/RAL_CLOSE_ maps calculated with CryoSPARC. Central slice (*right*) view is shown and colored according to the local resolution, as indicated in the scale bar. **i, j,** Orientation distribution sphere of the particles that contributed to the final CSC/RAL_OPEN_ and CSC/RAL_CLOSE_ cryo-EM reconstructions according to the relative signal, as indicated in the scale bar.

**Extended Data Figure 8.**
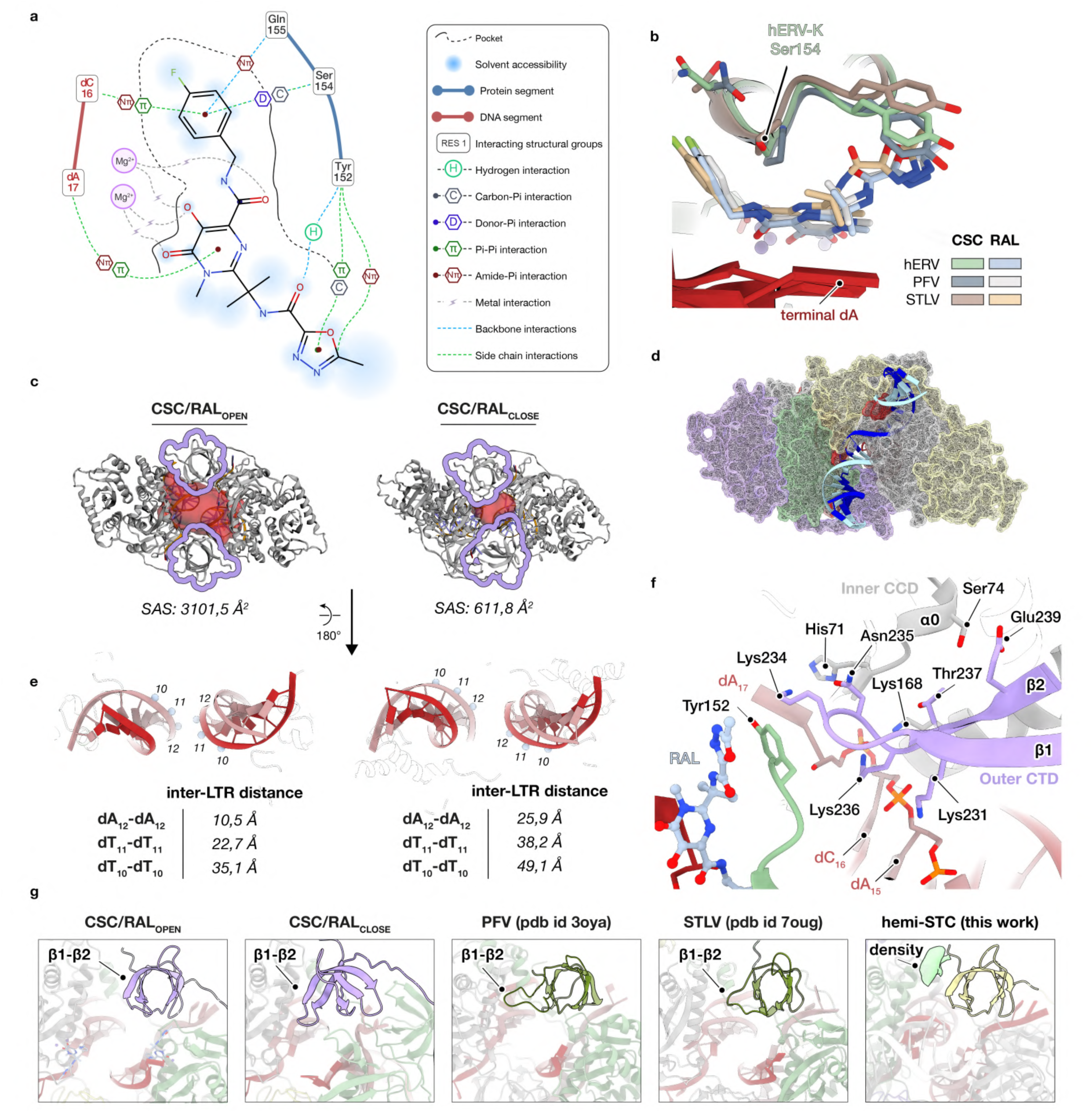
Structural characterization of Raltegravir (RAL) binding to hERV-K intasome. **a,** 2D diagram showing interacting residues, contact network and solvent accessibility surrounding Raltegravir (RAL) canonical binding sites in CSC/RAL_OPEN_ state. **b,** Structural comparison of RAL binding to hERV-K (this work, light green/light blue), PFV (pdb id 3oya, dark grey/light grey) and STLV (pdb id 7oug, brown/wheat) intasomes. Models are shown as ribbons and representative side chains are highlighted. Displaced integrating LTR strand is depicted in red. **c,** Surface topography analysis of hERV-K CSC/RAL_OPEN_ (*left*) and CSC/RAL_CLOSE_ (*right*) states on CASTpFold. The pocket surrounding the tDNA binding cleft is represented as a red density and solvent-accessible surface (SAS) values are indicated. Outer CTDs are delimited by a purple profile. **d,** Structural superimposition of CSC/RAL_CLOSE_ (surface color) and hERV-K STC (ribbon DNA) models. tDNA-binding cleft narrowing prevents DNA recruitment. **e,** Conformational changes associated to CSC/RAL_CLOSE_ formation involve inter-LTR broadening (red ribbons). The position of three representative backbone phosphates is marked by blue spheres and their connecting distances on CSC/RAL_OPEN_ (*left*) and CSC/RAL_CLOSE_ (*right*) is summarized in the inset table. **f,** Ribbon representation of CSC/RAL_CLOSE_ interaction network between hERV-K outer CTD (purple), inner CCD (grey), RAL (light blue) and LTR strand (red). Side chains of interacting residues are represented as sticks. **g,** Comparison of synaptic CTD position. The area occupied by the outer CTD in hERV-K CSC/RAL_CLOSE_ (purple ribbon) overlaps with the position of the extended β1-β2 loop in the PFV and STLV intasomes (both dark green). This region further coincides with the unassigned density (light green) observed in hERV-K hemi-STC (yellow) described in this study.

**Extended Data Figure 9.**
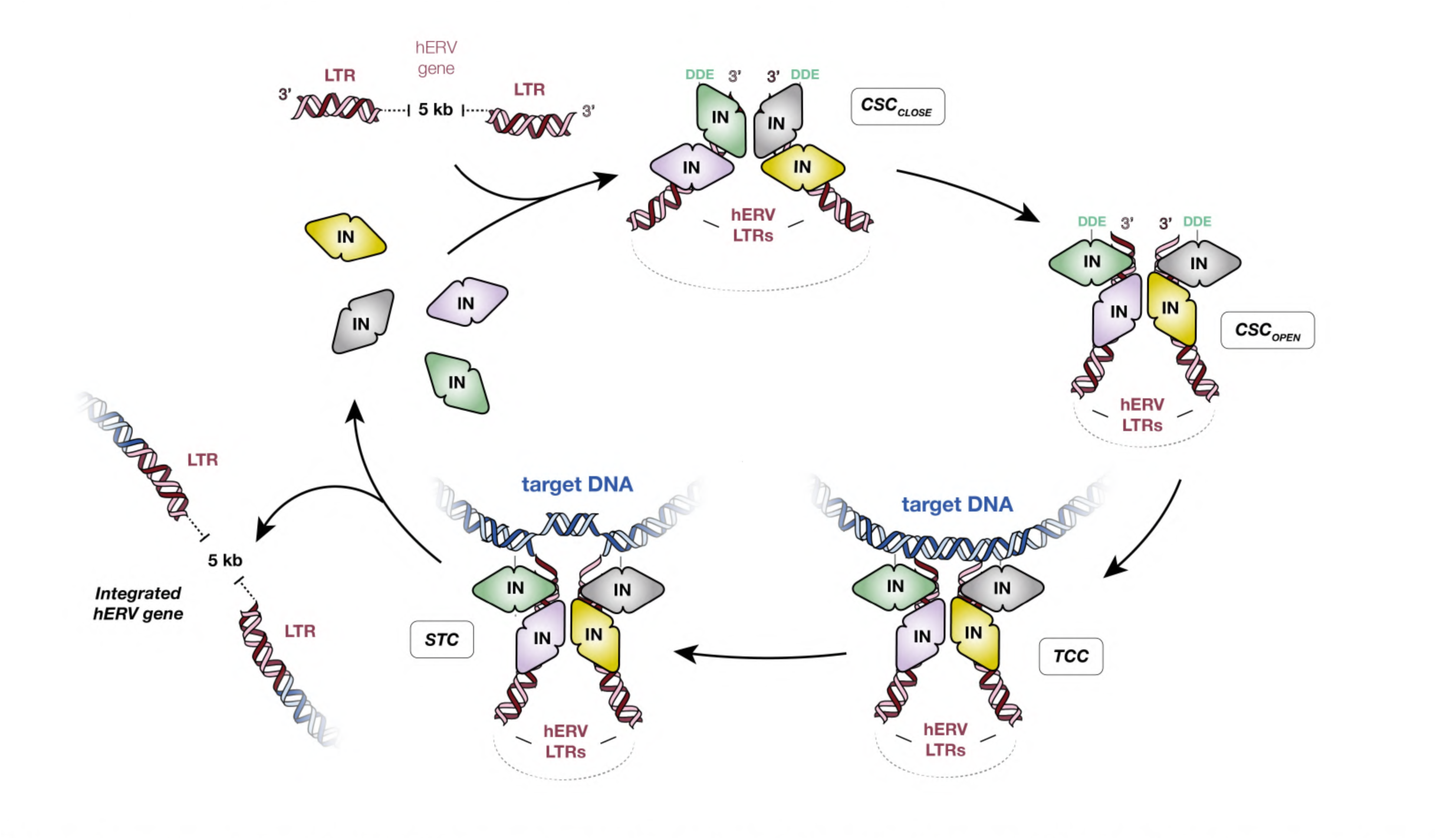
hERV-K intasome assembly and catalytic cycle Schematic representation of hERV-K intasome assembly and catalytic cycle. Integrase (IN) subunits are represented as geometrical shapes and colored as in Fig. 1a. The retrotransposon gene and target DNA (tDNA) are shown as red and blue helices, respectively. Following IN oligomerization around hERV-K long-terminal repeats (LTRs), the intasome complex transitions from a closed cleaved synaptic complex (CSC) state, which favors the accommodation of the gene termini but is deficient in tDNA binding, to an open CSC state, where the retrotransposon gene is stabilized and the tDNA-binding cleft is primed to target new genomic locations. This transitions into the target capture complex (TCC), which rapidly progresses to a strand transfer complex (STC) to culminate the integration reaction. Consequently, a copy of the retrotransposon is effectively mobilized, the complex disassembles, and the IN monomers are recycled for subsequent rounds of retrotransposition.

---

**Extended Data Table 1.**
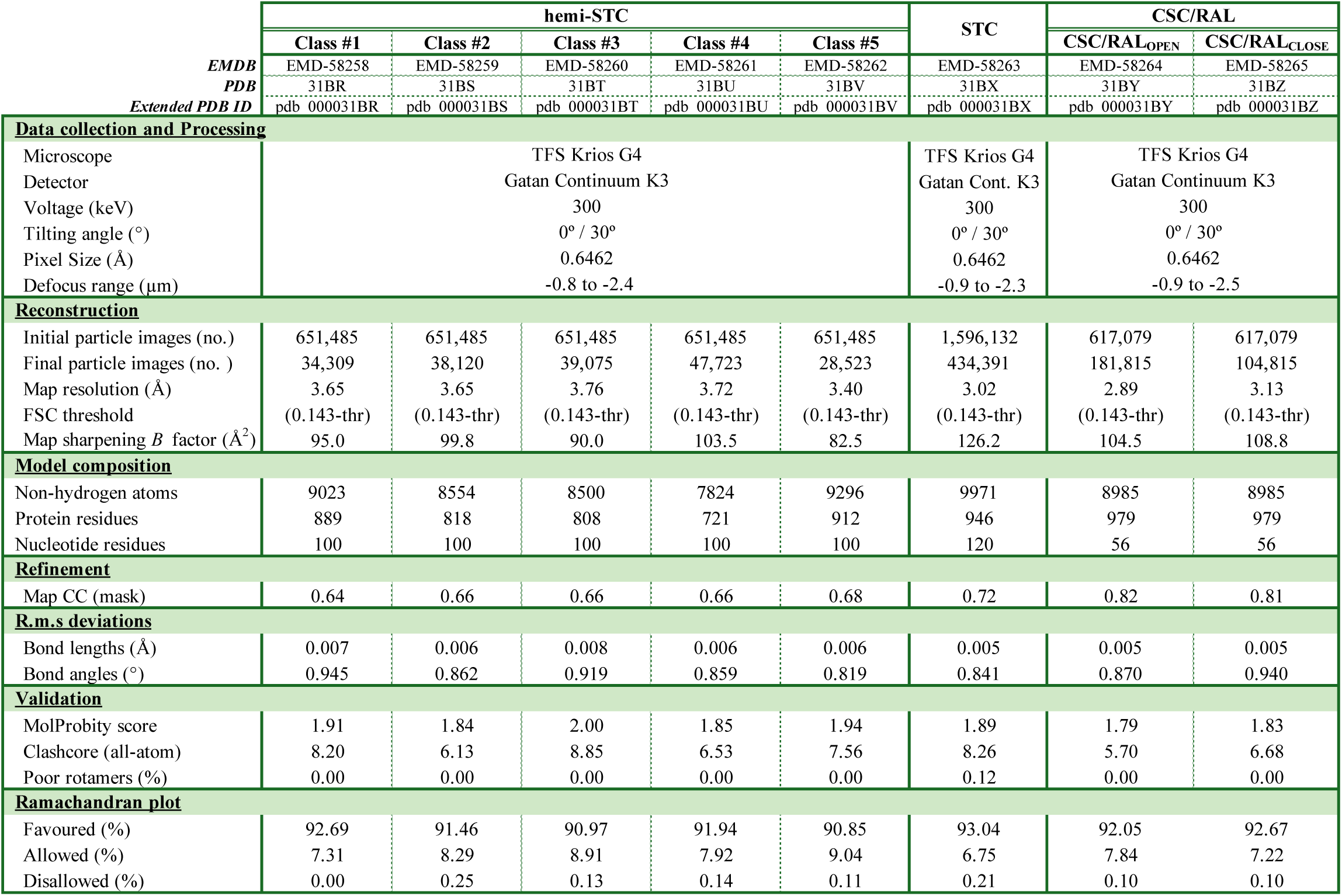
Cryo-EM data collection, refinement and validation statistics.

